# Cohesin loading at regulatory elements shapes 3D genome folding during erythropoiesis

**DOI:** 10.64898/2026.08.02.741886

**Authors:** Varshini Ramanathan, Chun-Jie Guo, Masahiro Nagano, Viraat Y. Goel, Clarice K.Y. Hong, Ariel D. Swett, Alexis Caulier, Emily King, Zuzana Tothova, Vijay G. Sankaran, Anders S. Hansen

**Author notes:** Co-first authors (Varshini Ramanathan, Chun-Jie Guo).

## Abstract

During erythropoiesis, differentiating cells silence much of the genome while maintaining high expression of select genes. It remains unclear whether this transcriptional specialization is linked to specific changes in 3D genome organization. We generated deep Micro-C maps across human erythropoiesis and identified erythroid *cis*-regulatory elements (CREs) that function as “matchmakers” by strengthening loops between neighboring elements. Matchmakers are associated with high H3K4me1, increase in strength during differentiation, and are linked to the expression of key erythropoiesis genes. Simulations suggest that a moderate cohesin loading bias across broad enhancer regions can explain the features of matchmakers, particularly under chromatin compaction. Targeted transcription factor and cohesin perturbations reduced matchmaker-associated structures and gene expression. We propose that matchmakers represent a subset of CREs that concentrate loop extrusion near key genes during erythroid differentiation.

## INTRODUCTION

During differentiation, cells convert broad developmental potential into specialized gene expression programs. Erythropoiesis, the specification of red blood cells (RBCs) from hematopoietic stem and progenitor cells (HSPCs), represents an extreme of gene regulatory specialization^1,2^. As progenitors mature into terminal erythroblasts, total RNA and protein content decline by ∼75%, yet hundreds of erythroid genes remain highly active or become strongly induced^3–8^. This creates a central regulatory problem: how do erythroid cells preserve and coordinate essential transcription while much of the genome is compacted and silenced?

Enhancers and other *cis*-regulatory elements (CREs) drive cell-type-specific transcription, often through enhancer-promoter (E-P) looping^9,10^. Cohesin mediates long-range E-P looping and proper expression of target genes, especially during cell-state transitions, though cohesin is not universally required for regulatory loops^11–21^. Cohesin forms loops and topologically associated domains (TADs) by loading onto chromatin and extruding until it dissociates or encounters convergent CTCF sites^22,23^. The distribution of cohesin across the genome has implications for control of cell-type specificity: if cohesin loads uniformly, then cell-type-specific genome structure is not influenced by the allocation of cohesin; if cohesin loads preferentially, then this spatial loading bias could shape cell-type-specific 3D genome structure.

To what extent cohesin exhibits preferential loading continues to be debated. Enhancer-bound Mediator complex was proposed to recruit the cohesin loader NIPBL and cohesin to form cell-type-specific E-P loops^24^although this model has been challenged^25^and antibody artifacts create false-positive NIPBL enrichment at CREs^26,27^. Engineered enhancer insertions have been shown to recruit cohesin and create sub-TADs^28–30^. Further, recent studies have identified a signature of targeted, or strongly preferential, cohesin loading in 3D contact maps: a stripe that originates from the loading element and extends orthogonally from the contact-map diagonal. “Plumes” were reported in mESCs after double depletion of CTCF and the cohesin unloader WAPL^31^; “jets” were reported in quiescent murine thymocytes^32^; “fountains” were reported to follow zygotic genome activation in zebrafish embryos^33^ and also found in *C. elegans*^34,35^. For simplicity, we will use the term “*fountain*” here. Aggregate fountain-like patterns have also been observed in T cell progenitors and subtypes^36–38^. However, single-locus observation of fountains has been exceedingly rare despite extensive 3D genome mapping across cell types and states. Furthermore, a recent study that exhaustively searched for signatures of cohesin loading at enhancers concluded that targeted loading contributes minimally to loop extrusion features in mESCs^39^. Indeed, artificially inducing targeted cohesin loading at enhancers inhibits target gene expression rather than enhancing it^39,40^. While this indicates that cohesin loading at an enhancer may not readily support long-range activation of its own target gene, whether targeted loading can instead mediate looping by neighboring enhancers remains unexplored.

Addressing this gap requires application of 3D genomics of sufficient resolution to resolve CRE looping and 3D structure at single loci. Here, we mapped 3D genome structure across human erythropoiesis using Micro-C. Unexpectedly, we identified a class of erythroid CREs that rewire local genome folding by promoting loops between flanking elements indirectly without engaging as a loop anchor directly. We call these elements matchmakers. Matchmakers are bound by key erythroid transcription factors (TFs), strengthen during differentiation, and are associated with the expression of essential erythroid genes. Polymer simulations, as well as TF and cohesin perturbations, support a model in which matchmakers preferentially load cohesin. We propose that matchmakers contribute to the regulatory specialization of erythropoiesis by concentrating cohesin-mediated genome folding at key erythroid regulatory loci.

## RESULTS

### Micro-C across human erythropoiesis reveals changes in genome organization across scales

To study the interplay of 3D genome structure and gene regulation during erythropoiesis, we induced differentiation of primary CD34^+^ HSPCs isolated from two independent human donors into mature erythrocytes as described previously^6^. We performed Micro-C^41^ at three timepoints corresponding to early, intermediate, and late erythropoiesis, which we refer to as Phase 1/P1, Phase 2/P2, and Phase 3/P3 (**Fig. 1A, S1A**). At ∼7B unique ligations per phase, this dataset represents the deepest 3D genomic data across erythropoiesis to our knowledge and provides unprecedented resolution (**Fig. S1B-E**).

**Figure 1:**
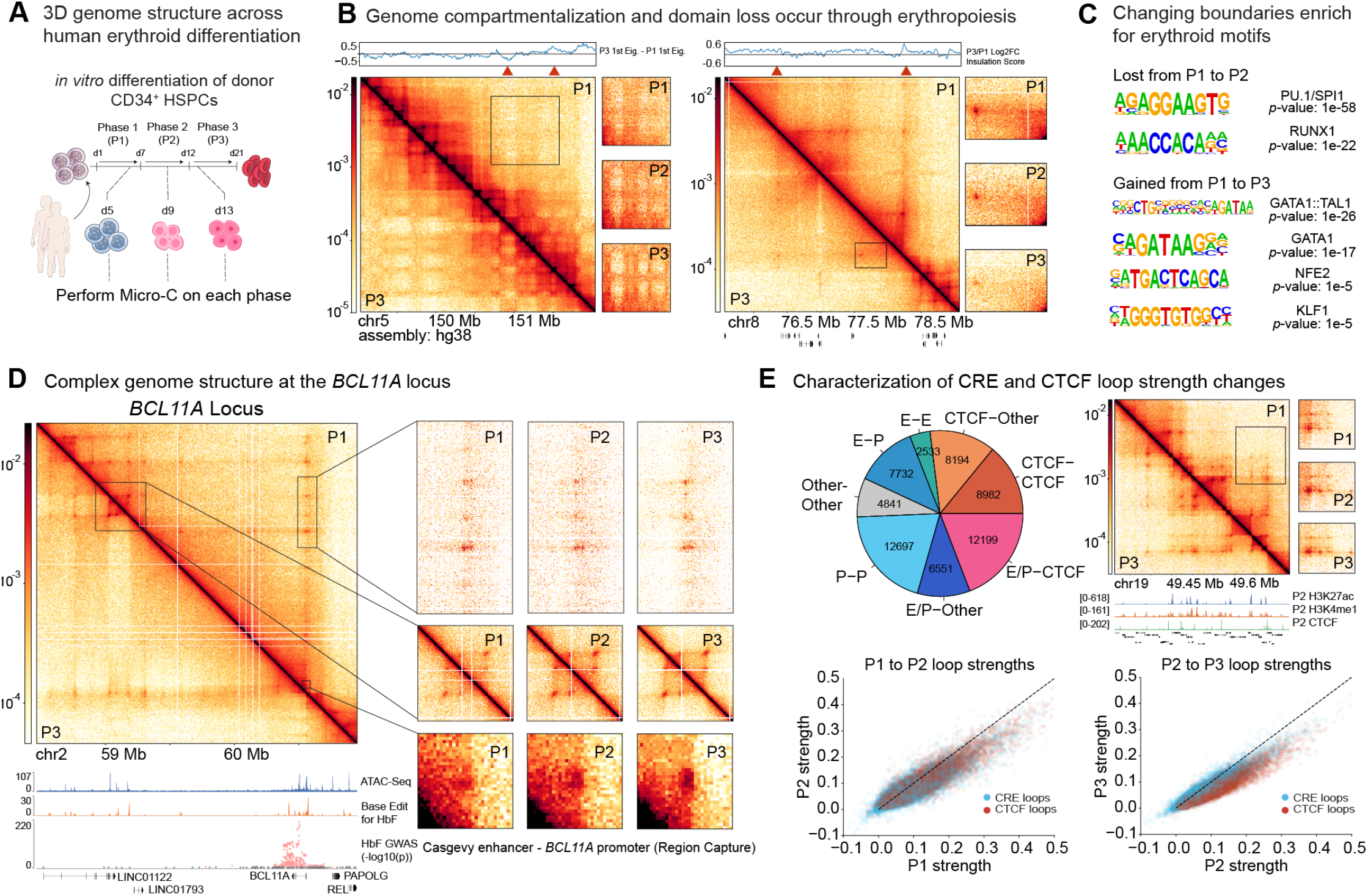
3D genomic mapping across human erythropoiesis. **(A)** Schematic representation of *in vitro* erythroid differentiation from primary human HSPCs. **(B)** Representative examples of genome compartmentalization (chr5:149023111-152003168 at 100kb binsize) and domain loss (chr8:75561811-79081548 at 10kb binsize) during erythropoiesis. Eigenvectors and insulation scores are computed with cooltools^80^ at the visualization binsize, with a 100kb window for insulation score. **(C)** Representative enriched motifs (called using HOMER) on changing boundaries, as identified with cooltools insulation. **(D)** The *BCL11A* locus (chr2:58464475-60946280 at 5kb binsize) over erythropoiesis. Bottom inset is Region Capture Micro-C (see Methods). Genomic tracks (from top to bottom): P1 ATAC-Seq^6^normalized base editing scores for HbF levels^48^and single-nucleotide polymorphisms associated with an increased HbF phenotype in a global genome-wide association study^81^. **(E)** (Top left) Annotations of 63,729 loops called by Mustache. (Top right) Representative example of CRE loops strengthening over erythropoiesis while a CTCF loop is maintained (locus: chr19:49314500-49733500 at 1kb binsize). (Bottom) Loop strengths across differentiation as measured by AbLE score^53^. CRE loops are E-E, E-P, P-P, and E/P-other loops, while CTCF loops are E-CTCF, P-CTCF, CTCF-CTCF, and CTCF-other loops.

At large scales, the Micro-C maps were consistent with previous 3D genomic studies of erythropoiesis^42–44^. We observed increased long-range interactions from P1 to P3 and strengthened B-compartment interactions (**Fig. 1B, S2A-C**), consistent with the 5-to 10-fold increase in chromatin compaction that characterizes terminal erythropoiesis^42,45^. Changes in insulation boundaries reflected erythroid specialization from P1 to P3: lost boundaries were enriched for TF motifs functioning in HSPCs (ETS and RUNX family) over all boundaries, while maintained or strengthened boundaries were associated with erythroid lineage-specific motifs (GATA1, GATA1::TAL1) **(Fig. 1C)** and upregulated genes (**Fig. S2D**).

Next, we took advantage of the uniquely high resolution of our Micro-C data to explore previously unresolvable 3D genome features. The +58-kb distal intronic enhancer of *BCL11A* is the target of the FDA-approved CRISPR therapeutic for sickle cell disease and β-thalassemia, exagamglogene autotemcel (Casgevy)^46,47^. We zoomed into fine-scale 3D interactions at the *BCL11A* locus, a key repressor of fetal hemoglobin (**Fig. 1D**). We resolved the Casgevy E-P loop between the +58-kb enhancer and *BCL11A* promoter and found it to increase markedly in strength during erythropoiesis, as expected (**Fig. 1D**). We additionally identified very long-range >1 Mb E-P loops to the *BCL11A* promoter that are associated with polymorphisms correlated with increased fetal hemoglobin expression^48^ (**Fig. 1D**), suggesting that additional distal enhancers beyond the validated intronic enhancer may contribute to *BCL11A* expression.

To systematically call loops genome-wide, we applied Mustache^49^ on the merged Micro-C data across phases. We identified 63,729 loops, which we classify as enhancer (H3K27ac/H3K4me1 peaks), promoter (TSS), CTCF (CTCF ChIP-Seq peaks with the CTCF motif), or other based on published datasets^42,50–52^ (**Fig. 1E, Methods)**. We quantified loop strengths across differentiation using AbLE^53^. CTCF loop strength decreased on average across differentiation, consistent with prior work reporting loss of TADs bound by CTCF during erythropoiesis^42,43^. However, although CTCF loops are generally substantially stronger than CRE loops^14,53,54^CRE loops were strikingly maintained relative to CTCF loops from P2 to P3 **(Fig. 1E, S3A-C)**. Given that chromatin compaction strongly increases during erythropoiesis, our observations are consistent with our previous study showing that chromatin compaction specifically strengthens affinity-mediated CRE loops without affecting extrusion-mediated structural CTCF loops^55^. Thus, erythropoiesis may represent a uniquely favorable environment for CRE loop formation.

### Erythroid lineage CREs exhibit distinct 3D genome structure

We next explored the relationship between CRE looping and expression dynamics during erythroid differentiation. We identified examples where CRE loops strengthened or weakened concordantly with gene activation or silencing (**Fig. S3D**), suggesting that erythroid lineage-specific CREs may form a distinct subset of strong, activating loops. To identify these elements, we used our recent Perturb-Multiome dataset^56^which defines genomic regions that change accessibility during erythropoiesis and/or following erythroid TF knockout (**Fig. 2A**). As expected, loops that overlapped TF-knockout differentially accessible (TF) and erythroid-specific (ery) regions were overwhelmingly CRE-CRE loops, which was especially true at the intersection of both sets (eryTF) **(Fig. 2A)**.

**Figure 2:**
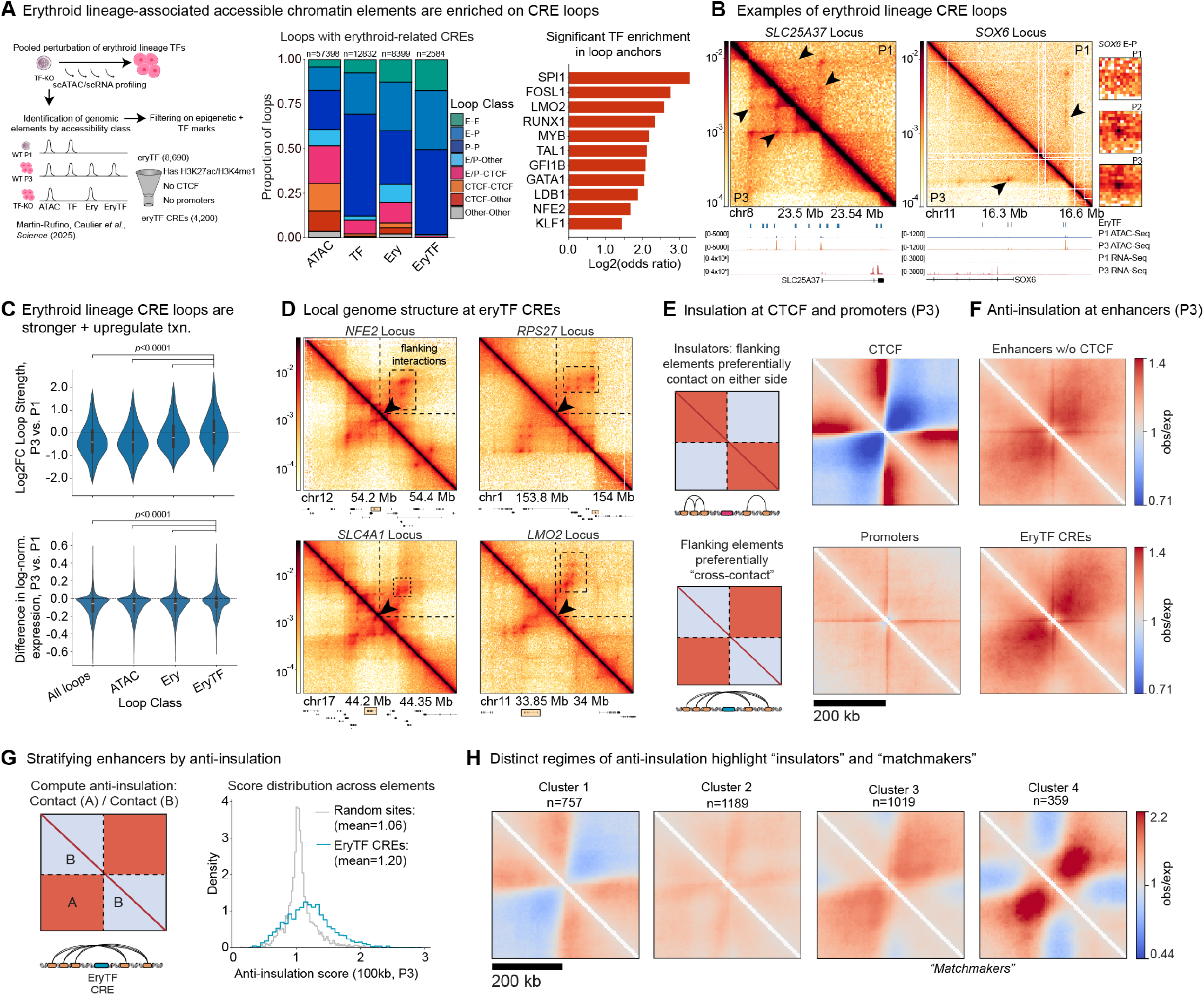
Genome structural features at erythroid lineage-specific elements. **(A)** (Left) Classification of erythroid-specific and TF-sensitive accessible chromatin regions from Martin-Rufino and Caulier *et al*.^56^followed by further filtering for putative erythroid-specific *cis*-regulatory elements (eryTF CREs). (Middle) Loop classifications of loops with both anchors occupied by ATAC, TF-sensitive (TF), erythroid-specific (ery), or erythroid-specific and TF-sensitive (eryTF) regions as annotated by Martin-Rufino and Caulier *et al*. (Right) Odds ratio enrichment of observing eryTF regions on loop anchors over observing ATAC peaks on loop anchors. Reported enrichments have p<0.0001 by Fisher’s exact test and adjusted FDR<0.01 by the Benjamini-Hochberg method. **(B)** Representative examples of eryTF regions on loop anchors at the *SLC25A37* (chr8:23458956-23585463 at 1kb binsize) and *SOX6* (chr11:16066449-16709591 at 2kb binsize; arrow indicates loop shown in inset) loci. Arrows indicate eryTF loops. **(C)** (Top) Changes in loop strength (AbLE score) at CRE loops (E-P/P-P/E-E) with erythroid-associated annotations on both anchors. Sample sizes: n=19743 for all loops, n=19474 for ATAC, n=9547 for ery, and n=2052 for eryTF. (Bottom) Changes in transcription at CRE loops with a promoter at one end. A single anchor per promoter was retained based on maximum overlap. Sample sizes: n=6634 for all loops, n=6554 for ATAC, n=3737 for ery, and n=761 for eryTF. In both plots, asterisks indicate p<0.0001 via Kruskal-Wallis multiple comparison test with post-hoc Mann-Whitney U tests (two-sided), FDR-corrected via Benjamini-Hochberg. **(D)** Local 3D genome structure surrounding eryTF CREs at the *NFE2* (chr12:54105812-54505812) (top left), *SLC4A1* (chr17:44081815-44481815) (bottom left), *RPS27* (chr1:153688754-154088754) (top right), and *LMO2* (chr11:33744908-34144908) (bottom right) loci in P3 at 2kb binsize. EryTF CREs are annotated with an arrow at the center. Box with highlight indicates the gene of interest. **(E)** Schematic description of insulation vs. anti-insulation, with demonstrative pileups of insulation at CTCF boundaries and accessible promoters. **(F)** Local pileups at 200kb flanking enhancers without CTCF, as well as eryTF CREs. **(G)** Anti-insulation score, as computed by the observed-over-expected contact of elements on either side of the scored element (region “A”) divided by the observed-over-expected contact of elements on the same side (region “B”). The score is computed at 100kb on either side of the central element. Shown are the anti-insulation score distributions for eryTF CREs deduplicated within 10kb (see Methods) and random sites (generated by shuffling ATAC intervals into the hg38 genome) sampled to the size of the deduplicated eryTF CRE distribution (3324), on P3 Micro-C. p=9.7×10^-77^ for the difference in means by the Mann-Whitney U Test (two-sided). **(H)** Pileups in P3 Micro-C of *k*=4 *k*-means clustering of eryTF CREs based on anti-insulation score across all three phases. Label indicates number of eryTF CREs (deduplicated within 10kb) in each cluster.

Additionally, loops were enriched for eryTF peaks over all ATAC peaks, suggesting that erythroid lineage-determining elements are preferentially engaged in loops **(Fig. 2A)**. For example, eryTF peaks form loops between the validated enhancers of *SLC25A37*^51^as well as a 227-kb E-P loop at *SOX6* **(Fig. 2B)**. EryTF CRE loops were stronger than typical CRE loops and eryTF-P loops were associated with higher gene expression changes than other CRE-P loops **(Fig. 2C, S3E-G)**.

Motivated by the key role of eryTF CREs, we explored the 3D genomic context surrounding these elements. Unexpectedly, we found that the strongest loops often formed between elements flanking the eryTF CRE, rather than with the eryTF CRE itself (**Fig. 2D**). Unlike canonical looping CREs, eryTF CREs appeared to promote loops between neighboring elements on either side. To globally investigate this, we performed pile-up analysis on all eryTF CREs as well as enhancers, CTCF sites, and promoters as comparisons. As expected, CTCF boundaries and promoters demonstrated insulation^57^: interactions are strongly reduced across CTCF sites and moderately reduced across active promoters (**Fig. 2E**). In contrast, CTCF-negative enhancers in general, and eryTF CREs in particular, showed the opposite pattern (**Fig. 2F**). Interactions that cross the eryTF CREs were substantially stronger than interactions on either side of the eryTF CREs, functionally amounting to “anti-insulation.”

We compared contacts that crossed each eryTF CRE (anti-insulation) with contacts confined to either side (insulation) within a 100-kb window, excluding CTCF-overlapping enhancers (**Fig. 2G, S4A, Methods**). Anti-insulation was significantly higher at eryTF CREs than at ATAC-Seq peaks without CTCF, accessible promoters, and at random genomic sites (**Fig. 2G, S4B-C**). We used *k-* means clustering to group eryTF CREs into 4 clusters **(Fig. 2H, S4D)**. Cluster 1 was broadly insulating, and was weakly enriched for CTCF (**Fig. S4E**), suggesting that these loci bind CTCF at levels that were undetected by our peak calling. Cluster 2 was approximately neutral, and clusters 3 and 4 exhibited anti-insulation in aggregate. We therefore refer to the 1,378 anti-insulating enhancers in clusters 3 and 4 as *matchmakers*, because they appear to promote looping between adjacent anchors on either side without themselves engaging in those loops as an anchor.

### Matchmakers can be explained by preferential cohesin loading and chromatin compaction

In aggregate, matchmakers resemble fountains^32,33,58^ (**Fig. 3A)**. However, whereas fountains appear as diffuse, loop-less stripes both in aggregate and at single loci, individual matchmaker loci are composed primarily of discrete loops rather than stripe-like patterns (**Fig. S5)**. To define this more rigorously, we divided loops into two categories: 1) “same-side loops” where both anchors are on one side of the matchmaker (akin to insulation) and 2) “crosser loops” where the loop anchors are on opposite sides of the matchmaker (akin to anti-insulation) (**Fig. 3A**). Crosser loop strength correlated strongly with anti-insulation intensity, whereas same-side loop strength did not (**Fig. 3A, S6A-B**). Crosser loops across matchmakers were also stronger than crosser loops across ATAC peaks or cluster 2 eryTF CREs, which lack aggregate anti-insulation, while same-side loop strengths (**Fig. S6C**) and loop-class distributions (**Fig. S6D**) were similar across groups. Thus, the distinguishing characteristic of matchmakers that drives anti-insulation is the strength of crosser loops. Matchmakers are not fountains at individual loci; instead, their aggregate fountain-like pattern is a “visual artifact” due to pileups aligning many discrete crosser loops of different sizes around the central matchmaker element.

**Figure 3:**
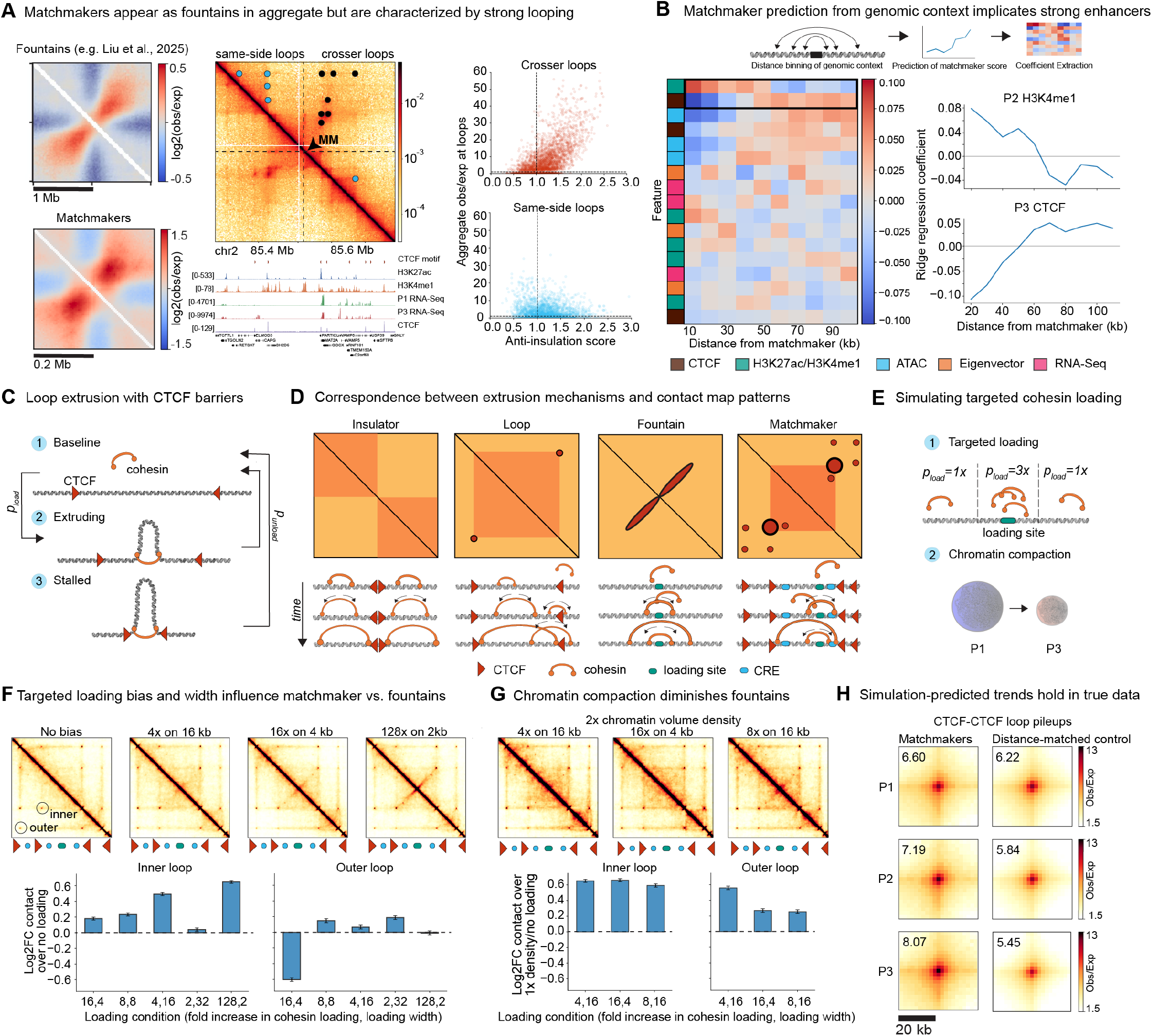
Moderate cohesin loading at enhancers can explain 3D genomic features of matchmakers. **(A)** Comparison of cluster 4 of matchmakers to fountains (Reproduced with permission under the Creative Commons Attribution 4.0 International License from: Liu, N.Q., Magnitov, M., Schijns, M.M.G.A., van Schaik, T., Teunissen, H., van Steensel, B., and de Wit, E. (2025). Extrusion fountains are restricted by WAPL-dependent cohesin release and CTCF barriers. Nucleic Acids Res. *53*. https://doi.org/10.1093/nar/gkaf549)^58^. However, individual matchmaker loci are composed of strong “crosser” loops rather than fountains (example locus: chr2:85303772-85703772 at 2kb binsize in P3 Micro-C; dashed line indicates eryTF CRE). **(B)** Feature coefficients, ordered by positive coefficient strength, of ridge regression to predict P3 anti-insulation score with inputs of P1-3 ATAC-Seq, RNA-Seq, CTCF, and H3K27ac, P1-2 H3K4me1, and P1-3 eigenvectors at 2kb as called by POSSUMM^82^. Inputs were binned by distance symmetrically from each eryTF CRE. **(C)** Schematic of cohesin-mediated loop extrusion with dynamic loading and unloading of cohesin. **(D)** Extrusion mechanisms yield distinct patterns on 3D genomic contact maps. **(E)** Schematic of simulation of targeted loading of cohesin with chromatin compaction. **(F)** Simulation of various targeted loading biases and loading widths (noted in kb; 1kb is one simulation monomer) in a 600kb locus inset with two convergent CTCF pairs, with quantifications for the outer pair and inner pair. **(G)** 2x chromatin volume density across different conditions of targeted loading. 4x bias on 16kb represents the ideal condition from (**F**), 16x bias on 4kb represents a condition which produced a fountain in (**F**), and 8x bias on 16kb tests the effect of increased bias on visual fountain appearance. **(H)** Comparison of pure CTCF-CTCF loops (with no other genomic annotations) within 100kb of matchmakers across differentiation (n=295) versus a distance-matched control of CTCF-CTCF loops (sampled to n=1000) with the same length distribution.

We next asked which genomic features predict anti-insulation. We used ridge regression to predict anti-insulation score from diverse (epi)genomic inputs (**Fig. 3B, S7A-C, Methods**). The model was moderately predictive (*R*2=0.43), indicating that matchmaker strength is partly encoded in local genomic context. The strongest positive predictors were proximal H3K4me1 and distal CTCF **(Fig. 3B, S7D)**. Consistently, strong matchmakers were enriched for broader and stronger H3K4me1 peaks relative to non-matchmaker eryTF CREs (**Fig. S8A**), and pileups stratified by H3K4me1 strength showed progressively stronger anti-insulation patterns (**Fig. S8B-C**). To determine whether this pattern was simply an artifact due to the position of matchmakers within TADs, we compared matchmakers to TAD-position-matched control sites. These controls showed that the strong anti-insulation observed at matchmakers is not explained by TAD position, either in aggregate (**Fig. S9**) or at single loci (**Fig. S10**).

Prior work has proposed that fountains arise from targeted cohesin loading, in which cohesin extrudes from a focal loading site to produce a stripe orthogonal to the diagonal on 3D maps^32,33,58^. We hypothesized that a moderate cohesin loading bias spread across several kilobases could strengthen crosser loops without producing visible single-locus fountains **(Fig. 3C-D)**. To test this, we performed 3D polymer simulations incorporating loop extrusion, CTCF boundaries, and affinity-mediated CRE-CRE interactions^53^ under varying cohesin loading regimes **(Fig. 3E, S11A)**. Indeed, strong focal loading produced visible fountains, whereas moderate but broad loading strengthened the nearest CTCF loop without generating fountain-like stripes (**Fig. 3F, S11B-C)**. Because erythropoiesis is accompanied by global chromatin compaction^42,45^we additionally simulated an increase in chromatin volume density under the same loading conditions. Strikingly, chromatin compaction reduced the visual prominence of fountains in contact maps. With only a twofold increase in chromatin volume density, crosser loops and domain strength were still increased by moderate loading biases, while diminishing the visual fountain-like appearance of targeted loading at stronger loading biases and smaller loading widths (**Fig. 3G, S11C-D**). Thus, preferential cohesin loading at matchmakers combined with chromatin compaction can strengthen local crosser loops without producing observable single-locus fountains.

Our polymer simulations predict that CTCF loops that form across matchmakers should strengthen during erythropoiesis, despite our observations that CTCF loops globally weaken during differentiation (**Fig. 1E**). To test this, we separated CTCF loops into matchmaker crosser loops and distance-matched non-matchmaker loops. Confirming our predictions, CTCF loops crossing matchmakers strengthened during differentiation and were strongest in P3, whereas matched loops not near matchmakers weakened (**Fig. 3H**). Consistently, cohesin occupancy as measured by RAD21 ChIP-Seq signal was enriched not only at matchmakers but also in the flanking regions, representing cohesin accumulation **(Fig. S12)**. Together, these results support a mechanistic model in which preferential cohesin loading at matchmakers under chromatin compaction is sufficient to explain the defining features of matchmaker genome structure: strengthened crosser loops, absence of single-locus fountains, and local loop strengthening despite global CTCF-loop weakening.

### Matchmakers are linked to erythroid gene expression and lineage TFs

Consistent with increased matchmaker-associated loop strengths during differentiation, matchmaker anti-insulation increased significantly during erythropoiesis **(Fig. 4A, S13A)**, with stronger matchmakers tending to increase the most (**Fig. S13B**). This differentiation specificity, as well as the association between matchmakers and erythroid enhancers **(Fig. 3B)**, suggests that cohesin loading at matchmakers could relate to the expression of erythroid-lineage genes. Using single-cell RNA-Seq, we identified P3 marker genes for late erythropoiesis and, as a negative control, marker genes for pooled non-erythroid progenitors (eosinophils, basophils, and mast cells)^56^. Indeed, P3 marker genes were significantly enriched for proximity (within 100 kb) to matchmakers over accessible promoters (**Fig. 4B**). This enrichment reflected erythroid specificity rather than transcriptional activity alone: P3 marker gene enrichment increased with matchmaker strength, whereas expression-comparable P3 genes were not more enriched in stronger matchmaker clusters (**Fig. S13C**). Genes near strong matchmakers were also more highly expressed in general (**Fig. 4C**). These associations may reflect cohesin extrusion from matchmakers strengthening loops between erythroid gene promoters and CREs on the other side of the matchmaker, which would result in stronger crosser loops. Indeed, crosser loops between promoters of marker genes and expressed genes were stronger than same-side promoter loops specifically at matchmakers (**Fig. 4D-E**). Thus, matchmakers are strongly associated with erythroid-specific gene expression.

**Figure 4:**
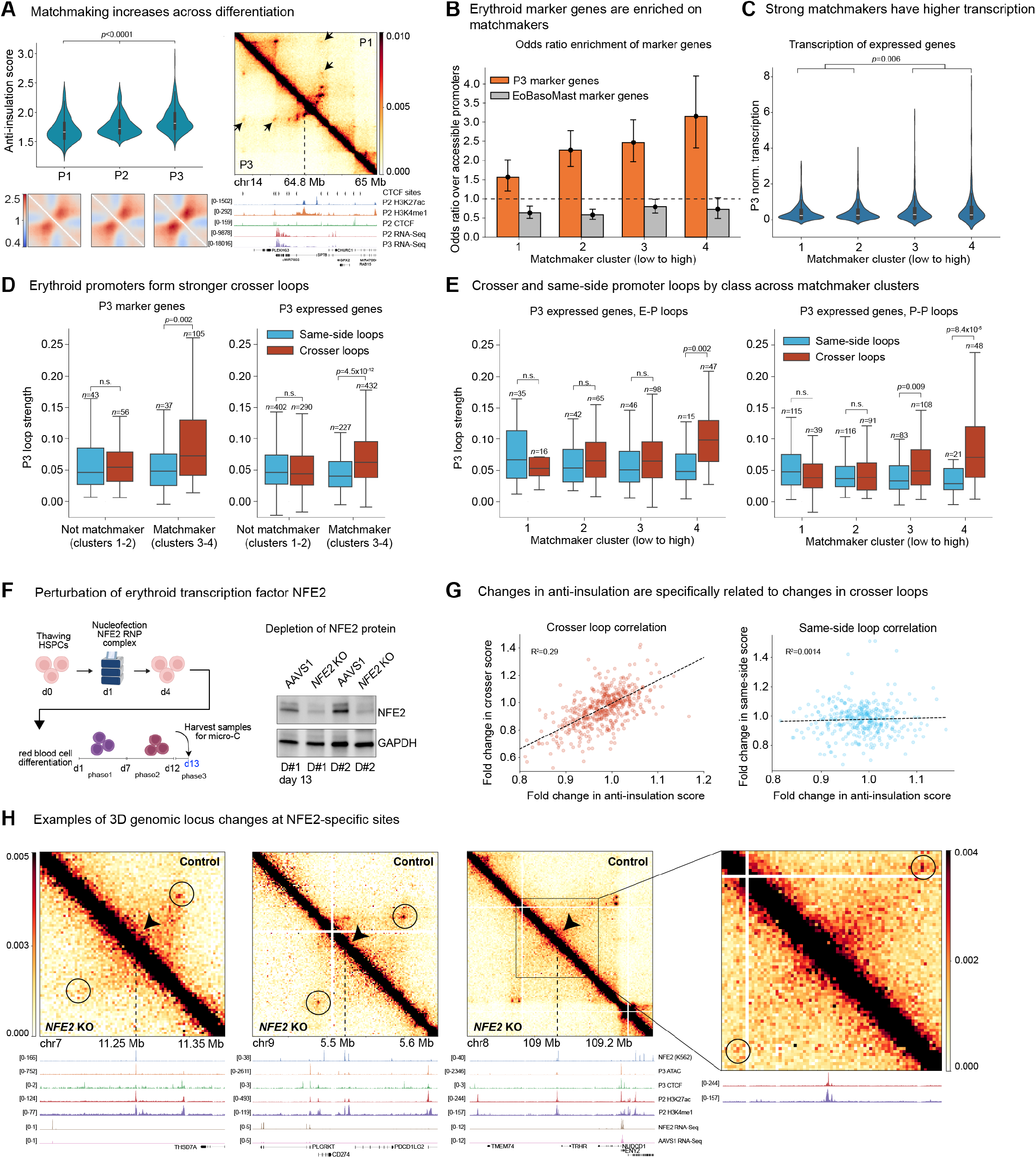
Matchmakers are functionally related to erythroid differentiation. **(A)** Anti-insulation score, and corresponding pileup, of Cluster 4 of matchmakers over differentiation (means across phases tested with a two-sided Wilcoxon signed-rank test between each pair) along with a representative example at the *SPTB* locus (chr14:64648114-65048114 at 2kb binsize). Arrows indicate crosser loops that increase over differentiation. **(B)** Odds ratio of observing each matchmaker cluster closest to marker genes corresponding to the P3 erythroid stage (polychromatic and orthochromatic erythroblasts) within 100kb, as compared to marker genes for diverging hematopoietic lineages (eosinophils, basophils, and mast cells). Error bars indicate 95% confidence interval. Clusters 1 and 2 are non-matchmaker eryTF CREs (i.e., not anti-insulating); clusters 3 and 4 are matchmakers. Odds ratios are computed using accessible promoters as the background set and statistically tested with Fisher’s exact test. n=72 for marker genes nearest cluster 1 (p=0.0008), n=125 for cluster 2 (p=1.12×10^-13)^n=103 for cluster 3 (p=1.18×10^-13^), and n=56 for cluster 4 (p=4.09×10^-12^). **(C)** Pseudo-bulked normalized transcription at each matchmaker cluster. p=0.006 for difference in means between matchmakers (cluster 3, n=633 genes and cluster 4, n=270 genes) and non-matchmaker eryTF CREs (cluster 1, n=916 genes and cluster 2, n=916 genes) by two-sided Mann-Whitney U test. Boxplots indicate median and IQR. **(D)** Comparison of P3 loop strengths (AbLE) of crosser loops and same-side loops to gene promoters near matchmaker (clusters 3 and 4) and non-matchmaker (clusters 1 and 2) eryTF CREs, identified as in **(B)** and Fig. S13C. *p*-value indicates the result of a two-sided Mann-Whitney U Test. Boxplots indicate median and IQR. **(E**) Comparisons as in **(D)** (right panel), for E-P and P-P loops specifically and split across matchmaker clusters. **(F)** Schematic and Western blot for knockout of *NFE2* followed by 13 days of erythroid differentiation to P3. D#1 and D#2 refer to independent donors. **(G)** Correlation between crosser loop fold change and anti-insulation score fold change (R^2^=0.29, n=491) versus between same-side loop fold change and anti-insulation score fold change (R^2^=0.0014, n=368) (*NFE2* KO/AAVS1 control). **(H)** Individual examples of changes in crosser loops where the NFE2-sensitive CRE (center of locus) is between loop anchors that do not overlap NFE2-sensitive CREs. From left to right, loci coordinates are chr7:11152122-11402122 (3.2kb binsize), chr9:5344067-5644067 (2kb binsize), and chr8:108702659-109472659 with inset chr8:108892659-109252659 (5kb binsize).

Because matchmakers are a subset of CREs bound by erythroid TFs, perturbing a lineage TF may disrupt matchmaker structure. We focused on NFE2 because it is not essential for erythropoiesis in mice^59^ and NFE2-perturbed primary human cells progress to late erythropoiesis^56^minimizing confounding effects from gross differentiation defects. We used CRISPR to knock out NFE2 in primary CD34^+^ HSPCs, induced erythroid differentiation, and performed Micro-C and RNA-Seq at P3 (**Fig. 4F**). Consistent with prior reports, NFE2 knockout had no detectable effect on erythroid maturation, with NFE2-perturbed cells showing CD71/CD235a profiles comparable to AAVS1 control cells (**Fig. S14A**).

To assess the effect of NFE2 perturbation on matchmakers, we repeated the anti-insulation score analysis in AAVS1 control and NFE2-KO Micro-C data, restricted to NFE2-sensitive TF-CREs. Anti-insulation decreased significantly, although modestly, in NFE2-KO cells relative to AAVS1 controls (**Fig. S14B-D**). The modest effect is likely explained by redundancy among erythroid TFs and incomplete editing after bulk nucleofection. Nevertheless, changes in anti-insulation score correlated specifically with changes in crosser loops (**Fig. 4G**), and individual loci showed weakened CTCF loops flanking strong NFE2 sites (**Fig. 4H**). CTCF crosser-loop intensity was lower in NFE2-KO cells than in AAVS1 controls compared with a distance-matched control set (**Fig. S14E**). Genes near matchmakers with reduced crosser-loop scores also showed lower transcription (**Fig. S14F**), suggesting a relationship between matchmaker-associated loop formation and gene expression. However, because NFE2 is itself a transcriptional activator, these experiments cannot fully separate changes in loops caused by alterations to loop extrusion from those caused by direct transcriptional effects.

### The matchmaker genome structure is dependent on cohesin

We next explored if matchmakers are cohesin dependent as indicated by our polymer simulations (**Fig. 3E-G**) and cohesin enrichment near matchmakers (**Fig. S12B**). However, cohesin is essential, and degron tagging is not feasible in primary human HSPCs, which cannot be propagated over extended periods or manipulated as individual clones (**Fig. 5A**). We therefore used two complementary approaches: perturbation analyses in related cell lines and partial knockout in primary HSPCs followed by differentiation to P3 and Micro-C.

**Figure 5:**
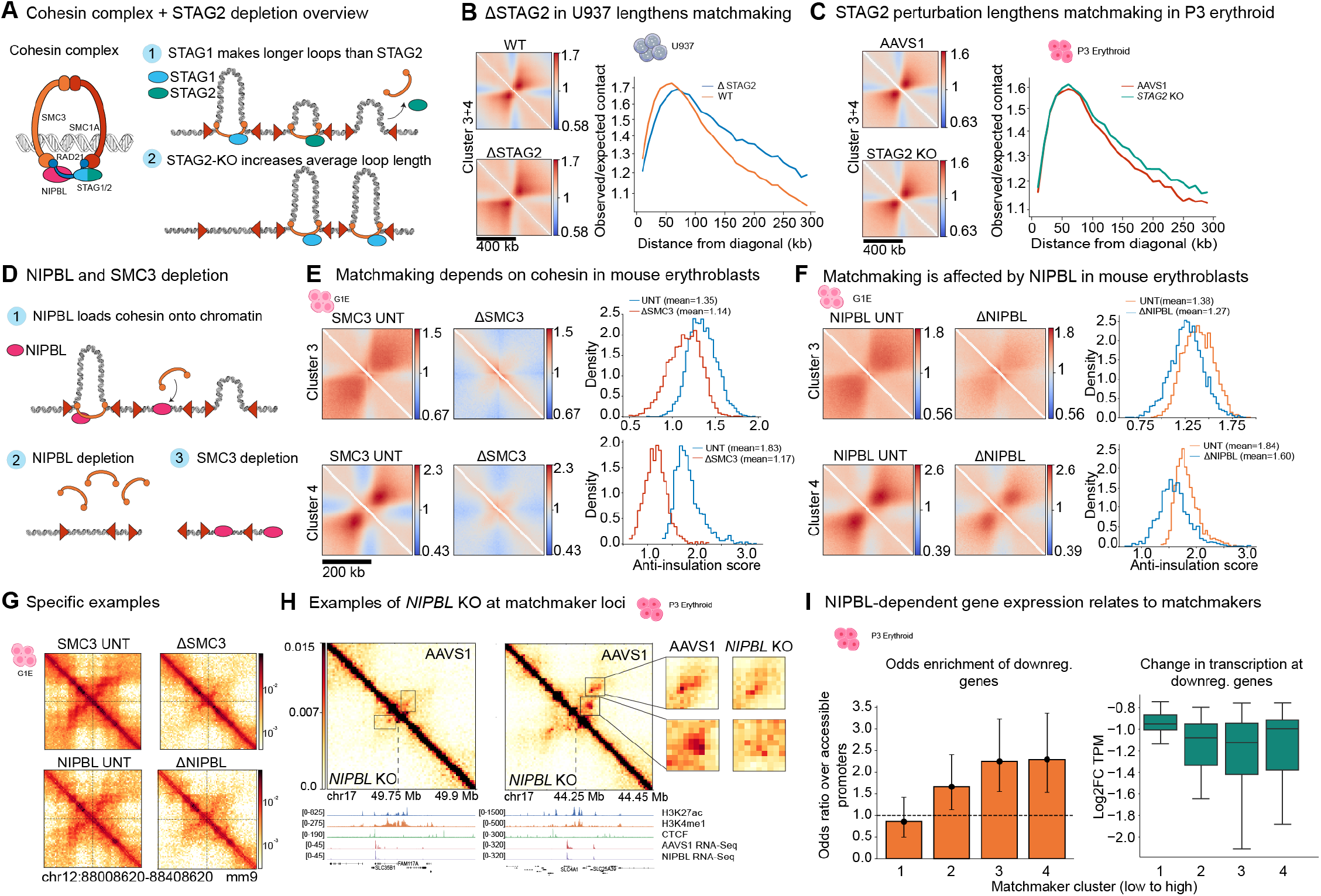
Cohesin perturbations affect matchmakers and associated gene regulation. **(A)** Diagram of the cohesin complex and the effect of STAG1 and STAG2 on cohesin loop length. **(B)** (Left) Pileups of candidate matchmakers identified in U937 monocytic cells. (Right) Quantification of diagonal extension by computing observed-over-expected contact on the axis orthogonal to the contact map diagonal in 10kb bins. (**C)** (Left) Pileups of matchmaker length extension in condition 1 of CRISPR knockout of *STAG2*. (Right) Quantification of STAG2 diagonal extension as in (**B**). **(D)** Schematic of expected effects of NIPBL and SMC3 perturbation. **(E)** Identification of matchmaker elements in G1E mouse erythroblast cells and pileup in SMC3 perturbation (Hi-C from Zhao et al., *Nature Genetics* (2024)^70^). Distributions describe anti-insulation score changes at Cluster 3 and 4 matchmakers . **(F)** Pileup of G1E matchmakers under NIPBL perturbation. Distributions describe anti-insulation score changes at Cluster 3 and 4 matchmakers. Hi-C from Aboreden and Zhao et al., 2026^11^. **(G)** Example locus (Hi-C visualized at 5kb binsize) demonstrating loss of crosser looping and anti-insulation in SMC3 and NIPBL depletion in G1Es. **(H)** Example loci (chr17:49588253-49958253 and chr17:44081815-44481815, visualized at 5kb binsize) where CTCF loops across matchmakers are perturbed upon *NIPBL* knockout in primary erythroid cells (condition 2). **(I)** (Left) Odds ratio of NIPBL-downregulated genes on matchmaker score clusters identified from all CTCF-negative enhancers. Error bars indicate 95% confidence interval. By Fisher’s exact test, p=0.86 for cluster 1, p=0.006 for cluster 2, p=2.13×10^-5^ for cluster 3, and p=2.13×10^-5^ for cluster 4. (Right) Extent of downregulation by Log2FC(TPM) of downregulated genes (averaged across conditions 1 and 2) at each cluster. In both, n=18 genes for cluster 1, n=42 for cluster 2, n=44 for cluster 3, and n=36 for cluster 4. Boxplots indicate median and IQR.

In somatic cells, the core cohesin complex comprises the subunits SMC1/3, RAD21, and STAG1/STAG2 (**Fig. 5A**). While the core SMC and RAD21 subunits are indispensable, STAG1 and STAG2 are partially redundant. STAG1 has a longer residence time and extrudes long loops, whereas STAG2 has a shorter residence time and extrudes shorter loops^60^. Thus, if matchmakers load cohesin, the anti-insulation pattern should expand in length away from the diagonal upon STAG2-KO because STAG1-cohesins will extrude longer on average (**Fig. 5A**). To test this, we performed Micro-C in the human monocytic cell line U937, comparing WT vs. STAG2-KO cells^61^ and identified matchmaker-like elements using the same anti-insulation scoring and clustering analysis as described previously **(Fig. 2G-H, Methods)**, excluding CTCF and promoter annotations from enhancer-marked regions^62^. As predicted, the intensity and length of matchmaking substantially increased in STAG2-KO U937 cells (**Fig. 5B**).

It is well-known that 3D genome architecture is remarkably robust to partial depletion of CTCF and cohesin^63–66^. Nevertheless, and anticipating much smaller effect sizes, we performed CRISPR knockout of STAG2 in primary HSPCs at day 0 (denoted “condition 1” or “Con1”) or at both days 0 and 2 (denoted “condition 2” or “Con2”) **(Fig. S15A-B, Methods)**. We then induced erythroid differentiation and performed Micro-C at P3. We analyzed all enhancers without CTCF, as opposed to just eryTF CREs, to be consistent with the U937 analysis (**Fig. 5B**). We similarly observed a modest increase in matchmaker-associated loop length in our primary erythroid cells upon partial STAG2 KO (**Fig. 5C, S15C**), consistent with the preferential cohesin loading model. STAG1 perturbation had little effect **(Fig. S15D)**, possibly because of compensation between STAG1 and STAG2 or a preference for STAG2-containing cohesin in erythroid cells^67–69^.

Having perturbed extrusion processivity, we disrupted extrusion altogether by depleting SMC3 (core subunit) and NIPBL (loading and extrusion factor) (**Fig. 5D)**. We re-analyzed Hi-C studies that acutely depleted SMC3^70^ and NIPBL^11^ in G1E mouse erythroblasts and we called G1E matchmakers as above (**Methods**) using G1E enhancer, promoter, and CTCF annotations^55,71^. Anti-insulation at matchmakers was almost completely lost upon SMC3 depletion, with the most pronounced effect at the strongest matchmakers (**Fig. 5E**). NIPBL depletion similarly diminished anti-insulation, albeit with a reduced effect size (**Fig. 5F-G**). Re-analysis of nascent transcriptomics (TT-Seq) upon 45 min of NIPBL depletion^11^ demonstrated that more downregulated genes (p-adj<0.05, log2FC<-0.5) were enriched near stronger matchmakers **(Fig. S15E)**.

Finally, we performed partial NIPBL knockout in primary human donor-derived HSPCs, followed by erythroid differentiation and Micro-C and RNA-Seq analysis at P3. Although editing was efficient, the aggregate effect on anti-insulation was modest (**Fig. S15F-G**). Nonetheless, the strongest NIPBL-sensitive matchmakers showed clear reduction in crosser loop strength at key erythroid genes, including *SLC4A1* (**Fig. 5H, Fig. S15H-I**). We identified 155 downregulated genes between the NIPBL-KO and AAVS1 control conditions (p-adj<0.05, log2FC<-0.5) and found these genes to be enriched near matchmakers with greater downregulation (**Fig. 5I**).

Taken together, these perturbation experiments of STAG2, SMC3, and NIPBL strongly implicate preferential cohesin loading at matchmakers as the causal factor for the anti-insulation genome structure pattern we observed and suggest a direct functional role for matchmaker CREs in controlling local gene expression during erythroid differentiation.

## DISCUSSION

During erythropoiesis, cells restrict global transcription while sustaining high expression of key erythroid genes. Here, using high-resolution Micro-C across primary human erythropoiesis, we identified matchmakers, a subset of erythroid CREs with a distinctive anti-insulation pattern. Rather than acting primarily as loop anchors, matchmakers strengthen loops between flanking CREs and CTCF sites. Matchmakers are associated with broad enhancer signatures, increase in strength during differentiation, are enriched near erythroid marker genes, and are sensitive to perturbation of erythroid transcription factors and cohesin machinery. Together with polymer simulations, these results support a model in which broad, moderate cohesin loading bias at matchmakers generates strong crosser loops and concentrates genome folding at loci that must remain actively transcribed throughout erythroid differentiation (**Fig. 6A**).

**Figure 6:**
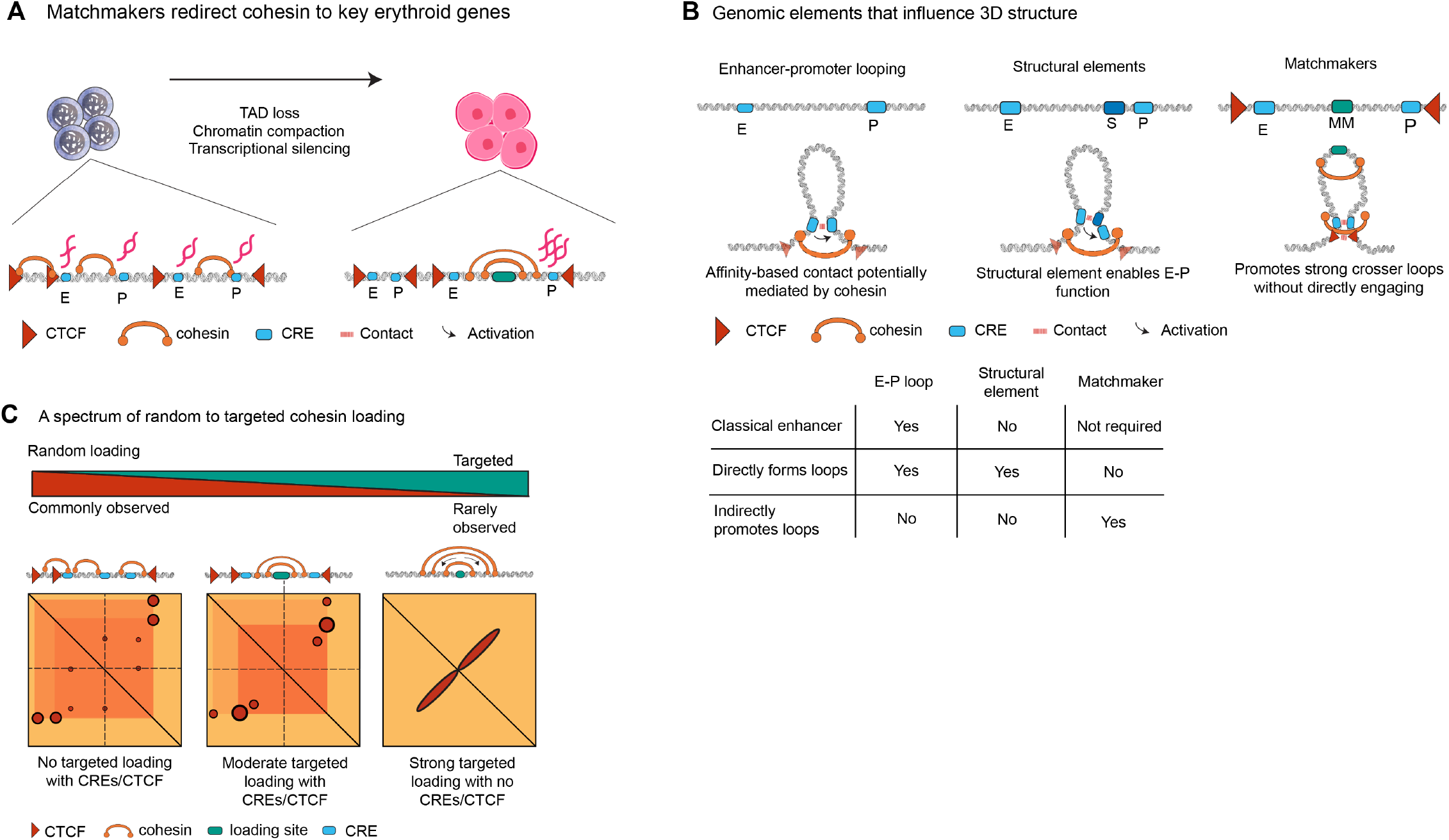
Proposed model of matchmaker-mediated cohesin loading during erythroid differentiation. **(A)** Proposed model by which matchmakers maintain genome organization and sustained gene expression at key erythroid loci by preferential cohesin loading. **(B)** Comparison to other named genomic elements that have been proposed to enact their effects via changes in 3D genome structure. **(C)** Schematic showing a progression from completely uniform to completely targeted loading and the expected effect on 3D genome structure.

This definition distinguishes matchmakers from other named CREs. Regulatory elements are generally defined by function or mechanism^72^. Transcriptional enhancers, range extenders, and facilitators are functionally defined: range extenders are required for certain E-P contacts to occur over long genomic distances^73^whereas facilitators are components of multipartite enhancers that have little classical enhancer activity alone but are required for full transcriptional activation of the gene loci in which they reside^74,75^. The mechanistic basis for facilitators and range extenders remains unclear^75^but they may operate as *structural elements* that form loops, thus bringing adjacent enhancers into proximity with target promoters (**Fig. 6B**). In contrast to functionally defined elements, we define matchmakers mechanistically: matchmakers are CREs that promote crosser loops between flanking CREs and CTCF sites, which our data suggest occurs through preferential cohesin loading at the matchmaker CRE **(Fig. 6A**). This further highlights a key distinction: matchmakers promote crosser loops largely indirectly, without serving as loop anchors, in contrast to CREs and structural elements that directly engage as a loop anchor (**Fig. 6B**).

Our findings support an intermediate model between uniform and targeted cohesin loading. Entirely uniform loading cannot readily explain why selected erythroid CREs strengthen loops across themselves, whereas strong focal loading as an exclusive loading mechanism would be expected to produce visible extrusion fountains at all enhancers. Matchmakers instead appear to reflect a broad, moderate cohesin loading bias at a subset of erythroid CREs (**Fig. 6C**). Polymer simulations show that this regime, especially when combined with erythroid chromatin compaction, strengthens crosser loops without producing single-locus fountains. Potential molecular bases of preferential cohesin loading at matchmakers include recruitment of NIPBL by transcription factors or transcriptional machinery^21,24,76^preferential loading in accessible enhancer chromatin, and enhancer-transcription-induced DNA supercoiling, because SMC complexes preferentially load on supercoiled DNA^77^.

Matchmakers are associated with erythroid gene regulation. They are enriched near P3 erythroid marker genes, and this enrichment reflects erythroid specificity rather than expression level alone. Genes near strong matchmakers are more highly expressed and NIPBL-sensitive genes are enriched near matchmakers. These results support a functional link between matchmaker-associated folding and the transcriptional specialization of erythropoiesis, although individual matchmaker deletions will be needed to establish direct regulatory requirements at specific loci.

Several limitations should be noted. It is challenging to study cohesin loading in endogenous contexts genome-wide since cohesin extrudes quickly away from loading sites to enrich at barriers^78^. Therefore, instead of directly observing single loading events, we infer preferential cohesin loading from contact maps, simulations, cohesin enrichment, and perturbations. The studies in primary human erythroid cells also limited complete cohesin depletion, degron tagging, and systematic genome editing. Finally, comparison to prior studies observing aggregate fountains is challenging due to their lower resolution^33,36–38^. While our high-depth Micro-C maps allowed us to resolve discrete single-locus crosser loops within previously reported aggregate anti-insulation or fountain-like pileups, similar investigations will be required to determine whether these conclusions generalize to other cell types.

Despite these limitations, matchmakers reveal a route by which lineage-specific CREs can shape genome architecture during differentiation. Rather than acting only as classical transcriptional enhancers, matchmakers bias cohesin-mediated extrusion toward loci that require sustained high-level transcription. Recent work has demonstrated that only longer-lived E-P contacts are transcriptionally productive^20^. Because cohesin extrusion is expected to stabilize E-P contacts and increase their duration, cohesin may serve a particularly important role in sustaining strong transcription during terminal erythropoiesis^13,20,79^. Future high-resolution Micro-C studies will reveal the generality of matchmaking in other developmental systems. We propose that matchmakers may represent a general mechanism by which cell-type-specific CREs instruct the 3D genome, linking lineage-specific gene expression to the spatial allocation of loop extrusion.

## Acknowledgements

We thank Henry Lu, Liam Cato, and Uma Arora for early contributions to the project including RCMC probe design and public dataset selection. We thank Nicholas Aboreden for assistance with G1E datasets. We thank Elzo de Wit, Gerd Blobel, Elphège Nora, and Erika Anderson for feedback on the manuscript. We thank Leonid Mirny and Aleksandra Galitsyna for discussion and feedback on the project. We thank the Hansen and Sankaran Labs, including Jamie Drayton, James Jusuf, Sarah Nemsick, and Jin Yang, for helpful discussions throughout the project and feedback on the manuscript. We thank Edward Banigan, James Jusuf, and Jin Yang for advice on polymer simulations. AI disclosure: AI tools were used as a grammatical assistant and minimal coding aid (code fragments for data organization and visualization). All resulting text was reviewed and further edited by humans.

## Author Contributions

V.R., C-J.G., A.S.H., and V.G.S. contributed to project design. C-J.G. and V.R. designed erythroid perturbation experiments with input from A.C., A.S.H., and V.G.S. C-J.G. performed and characterized erythroid differentiation, prepared cells for Micro-C, and performed and characterized genome editing experiments. A.D.S. performed U937 experiments with supervision from Z.T. V.Y.G. performed differentiation Micro-C and RCMC, M.N. performed erythroid perturbation Micro-C, and C.K.Y.H. performed U937 Micro-C. V.R. performed all analysis and interpretation including bioinformatics, Micro-C analyses, and polymer simulations. A.C. and E.K. prepared Perturb-Multiome datasets and provided input on analysis. V.R. prepared all figures. V.R. and A.S.H. drafted the manuscript with input from V.G.S. and with contributions to the methods from C-J.G., V.Y.G., and A.D.S. All authors edited the manuscript. A.S.H. and V.G.S. supervised the project.

## Data and Code Availability

Data generated for this study (Micro-C and RNA-Seq) can be found on GEO under accession numbers GSE341565, GSE341414, GSE341752 and GSE341753. Publicly available datasets re-analyzed for this study can be found in **Table S2**. Code for analyses and figure generation can be found on GitHub at: https://github.com/ahansenlab/erythroid_3D_genome.

## Materials and Methods

Materials and methods can be found in the Supplementary Information.

