## Supplementary Information for "Cohesin loading at regulatory elements shapes 3D genome folding during erythropoiesis"

### Supplementary Information for
### Cohesin loading at regulatory elements shapes 3D genome folding during erythropoiesis

Varshini Ramanathan[1-4, ^], Chun-Jie Guo[2,5,6,7,^], Masahiro Nagano[1-4], Viraat Y. Goel[1-4], Clarice K.Y. Hong[1-4], Ariel D. Swett[8,9], Alexis Caulier[2,5,6,7], Emily King[2,5,6,7], Zuzana Tothova[8,10], Vijay G. Sankaran[2,5,6,7,11,*] Anders S. Hansen[1-4,*]

[1]Department of Biological Engineering, Massachusetts Institute of Technology, Cambridge, MA 02139, USA; [2]Gene Regulation Observatory, Broad Institute of MIT and Harvard, Cambridge, MA 02139, USA; [3]The Novo Nordisk Foundation Center for Genomic Mechanisms of Disease, Broad Institute of MIT and Harvard, Cambridge, MA 02142, USA; [4]Koch Institute for Integrative Cancer Research, Cambridge, MA 02139, USA; [5]Division of Hematology/Oncology, Boston Children's Hospital, Harvard Medical School, Boston, MA 02115, USA; [6]Department of Pediatric Oncology, Dana-Farber Cancer Institute, Harvard Medical School, Boston, MA 02115, USA; [7]Howard Hughes Medical Institute, Boston, MA 02115, USA; [8]Department of Medical Oncology, Dana-Farber Cancer Institute, Harvard Medical School, Boston, MA 02115, USA; [9]Chemical Biology Program, Harvard University, Cambridge, MA 02139, USA; [10]Cancer Program, The Broad Institute of MIT and Harvard, Cambridge, MA 02139, USA; [11]Harvard Stem Cell Institute; Cambridge, MA 02138, USA; ^Co-first authors, *Co-corresponding authors:

### List of Supplementary Tables

**Supplementary Tables (attached):**
*Table S1:* Micro-C mapping information for all datasets.
*Table S2*: Public datasets used in this study.
*Table S3*: Guides and primers used in this study.

### Methods

#### Experimental Methods

##### Human primary HSPC culture and in vitro erythroid differentiation

Human $CD34^+$ hematopoietic stem and progenitor cells (HSPCs) isolated from mobilized peripheral blood of healthy adult donors were obtained from the Cooperative Center of Excellence in Hematology at the Fred Hutchinson Cancer Research Center. Cryopreserved HSPCs were thawed and cultured in maintenance medium consisting of StemSpan SFEM II (StemCell Technologies) supplemented with CC100 cytokine cocktail (StemCell Technologies), 50 ng/mL human thrombopoietin (TPO; PeproTech), 1% penicillin-streptomycin (Life Technologies), and 1% L-glutamine (Life Technologies).

Following the maintenance phase, erythroid differentiation was performed using a previously described three-phase culture system (1). Briefly, a basal erythroid medium was prepared using IMDM (Gibco) supplemented with 2% human AB plasma (SeraCare), 3% human AB serum (Life Technologies), 3 U/mL heparin, 10 μg/mL insulin, 200 μg/mL holo-transferrin, and 1% penicillin-streptomycin. During phase I (days 1–7), the basal medium was supplemented with 3 U/mL erythropoietin (EPO), 10 ng/mL human stem cell factor (SCF), and 1 ng/mL interleukin-3 (IL-3). During phase II (days 7–12), cells were cultured in basal medium supplemented with 3 U/mL EPO and 10 ng/mL SCF. During phase III (day 12 onward), cells were cultured in basal medium supplemented with 3 U/mL EPO and 1 mg/mL holo-transferrin.

##### Genome editing of human primary HSPCs

Genome editing was performed in primary human $CD34^+$ hematopoietic stem and progenitor cells (HSPCs) using the 4D-Nucleofector system (Lonza) and the P3 Primary Cell 4D-Nucleofector X Kit S (Lonza). For NFE2 knockout, ribonucleoprotein (RNP) complexes were assembled by combining 100 pmol Cas9 protein (IDT) with 100 pmol chemically modified sgRNA (Synthego) and incubating at room temperature for 15 min. RNP complexes were delivered into $CD34^+$ HSPCs using the P3 nucleofection reagent supplemented with Alt-R Cas9 Electroporation Enhancer (IDT) at a 20:1 molar ratio, using program DZ-100. For knockout of cohesin complex components (STAG1, STAG2, and NIPBL), two editing conditions were used to achieve different levels of gene disruption. In condition 1 (Con1), cells were edited using the same procedure described above. In condition 2 (Con2), cells underwent a second round of nucleofection 2 days after the initial editing using the same RNP delivery protocol. Following editing, cells were cultured under erythroid differentiation conditions beginning 3 days after the initial nucleofection. Knockout efficiency was assessed by western blot analysis of protein lysates or by genomic DNA extraction followed by ICE (Inference of CRISPR Edits) analysis.

##### Erythroid Micro-C sample preparation

Micro-C was performed on erythroid cells differentiated *in vitro* from human $CD34^+$ HSPCs. For erythroid differentiation time-course experiments, cells were collected on days 5, 9, and 13 from two independent healthy donors. NFE2-knockout samples were collected on day 13 from two independent healthy donors. STAG1-, STAG2-, and NIPBL-knockout samples were collected on day 13 from a single donor under two editing conditions. Cells were sequentially crosslinked with 3 mM DSG at a density of $1x10^6$ cells/mL for 35 min, followed by 1% formaldehyde for 10 min at room temperature. The crosslinking reaction was quenched with 0.375 M Tris-HCl (pH 7.5). The pellets were snap frozen in liquid nitrogen and stored at -80 °C for Micro-C.

##### U937 cell culture and Micro-C sample preparation

Isogenic U937 single-cell clones with and without STAG2 knockout were used, generated as described in (2). Three WT (Gc1a1, Gc3e1, Gc3e3) and two STAG2-KO (A2f11, A2g4) clones were thawed and cultured in RPMI (Thermo Fisher #11875119), 10% fetal bovine serum (Thermo Fisher #A5670701), and 1% Pen/Strep with glutamine (Life Technologies Corp 10-378-016). 30 million viable cells per genotype were collected (800g, 5min, room temperature), washed with PBS, and recollected. Cells were resuspended in Pierce DSG cross linker (Thermo Fisher #20593) at $1x10^6$ cells/mL and rotated for 35 minutes at room temperature. Formaldehyde was added dropwise to a final concentration of 1%, and cells were fixed for 10 minutes rotating at room temperature. Tris pH 7.5 was again added dropwise to a final concentration of 0.375 M, and cells were quenched for 5 minutes rotating at room temperature. The cells were then collected, washed with cold PBS twice, and aliquoted into 5 million cell pellets. The pellets were snap frozen in liquid nitrogen and stored at -80 °C for Micro-C.

##### Micro-C and Region Capture Micro-C

Micro-C and Region Capture Micro-C (RCMC) were performed largely according to Huseyin *et al.*, *Nature Protocols* (2026) (3) with minor modifications.

*Micrococcal nuclease (MNase) titration*
Ideal digestion conditions were determined for each batch of crosslinked cells by treating 1M cell samples with varying amounts of MNase and digesting at 37°C for 20 min on a thermomixer. Digested chromatin underwent crosslink reversal, DNA purification, and gel-based separation to visualize the fragment size distribution. Ideal digestion concentrations were identified by samples digested to primarily (~80%) monomeric fragments (150-200 bp), few (~15-20%) dimeric fragments (250-350 bp), and a faint but visible band (<5%) of trimeric fragments (400-500 bp).

*Micrococcal nuclease digestion*
Cell membranes were solubilized to extract intact nuclei by resuspending crosslinked cell pellets in Micro-C Buffer #1 (MB#1; 50 mM NaCl, 10 mM Tris-HCl pH = 7.5, 5 mM MgCl2, 1 mM CaCl2, 0.2% NP-40 Alternative (Millipore Sigma #492018), 1x Protease Inhibitor Cocktail (Sigma-Aldrich #5056489001)) at 1M cells per 100 µL for 20 min on ice. Following an MB#1 wash, samples were resuspended in 100 µL MB#1 and the ideal amount of 20 U/µL MNase (Worthington Biochem #LS004798) determined by the MNase titration was added. This digestion reaction was mixed at 37°C for 20 min on a thermomixer before being quenched with 4 mM EGTA (bioWORLD #40520008) and heat inactivated at 65°C for 10 min.

Digested nuclei were washed twice with ice-cold Micro-C Buffer #2 (50 mM NaCl, 10 mM Tris-HCl pH = 7.5, 10 mM MgCl2, 100 μg/mL BSA (Sigma-Aldrich #B8667)).

*End repair and labeling*

Digested fragments were enzymatically blunted and biotinylated. First, digested chromatin was 5' phosphorylated in end-repair reactions (50 U T4 Polynucleotide Kinase (New England BioLabs #M0201), 50 mM NaCl, 10 mM Tris-HCl pH = 7.5, 10 mM MgCl2, 100 ug/mL BSA, 2 mM ATP (Thermo Fisher #R1441), 5 mM DTT (Sigma-Aldrich #10197777001), in water) at 37°C for 15 min while mixing. Next, 50 U of DNA Polymerase I Klenow Fragment (New England BioLabs #M0210) was added to the reaction and incubated at 37°C for 15 min while mixing to create 5' fragment overhangs, and these overhangs were filled in by adding a mixture of dNTPs in end-labelling buffer (66 μM each of dTTP (Jena Bioscience #NU-1004), dGTP (Jena Bioscience #NU-1003), biotin-dATP (Jena Bioscience #NU-835-BIO14), and biotin-dCTP (Jena Bioscience #NU-809-BIOX), 1X T4 DNA Ligase Buffer, 100 μg/mL BSA, in water) and incubating at room temperature for 45 min with interval mixing. This end-blunting reaction was quenched by 30 mM EDTA (Invitrogen #15575020) and heat inactivated at 65°C for 20 min. Finally, end-blunted and biotin-labeled nuclei were washed once with Micro-C Buffer #3 (50 mM Tris-HCl pH = 7.5, 10 mM MgCl2, 100 μg/mL BSA).

*Proximity ligation and removal of unligated biotin*

Proximity ligation was performed by incubating labeled chromatin in a ligation reaction (10,000 U T4 DNA Ligase (New England BioLabs #M0202), 1X T4 DNA Ligase Buffer, 100 μg/mL BSA, in 500 μL water) at room temperature for at least 2.5 hours or overnight with gentle mixing. To remove biotinylated dNTPs from all unligated fragment ends, samples were digested by 1,000 U of Exonuclease III (New England BioLabs #M0206) in reaction buffer (1X NEBuffer #1 in water) at 37°C for 15 min with interval mixing.

*DNA purification and size-selection*

Ligated DNA fragments were purified over a series of steps. DNA was first reverse crosslinked to remove proteins and RNA by adding 1% SDS (Sigma-Aldrich #L3771), 2 mg/mL Proteinase K (Viagen Biotech #501-PK), 250 mM NaCl, and 100 μg/mL RNase A (Thermo Fisher #EN0531) to the samples and incubating at 65°C overnight. Following crosslink reversal, the DNA solution was purified using the Zymo DNA Clean & Concentrator kit (Zymo Research #D4034) according to the kit manual.

Ligated DNA fragments were subsequently size-selected (~200-400 bp) by extraction from a 1% agarose gel (VWR #97062). Gel extracts were purified using the Zymo Gel Purification kit (Zymo Research #D4008), and samples were quantified by Qubit 1X dsDNA High Sensitivity Assay (Invitrogen #Q33231).

Ligated fragments were further purified by using Dynabeads MyOne Streptavidin T1 (Invitrogen #65601) to enrich for biotinylated fragments. DNA samples were bound to beads in a Binding and Wash Buffer (1 M NaCl, 5 mM Tris-HCl pH = 7.5, 500 μM EDTA, 0.1% Tween-20 (Sigma-Aldrich #P8074)) at room temperature for at least 30 minutes with mixing. After two washes with the Binding and Wash Buffer, the bead-bound samples were washed once with 10 mM Tris-HCl pH = 7.5 prior to library preparation.

*Library preparation*

Illumina library preparation was performed using the NEBNext Ultra II kit (New England BioLabs #E7645), with the addition of interval shaking (1 minute on, 3 minutes off) at 1000 rpm during incubations to mix the bead-bound samples. Sample washes were performed using Binding and Wash Buffer and 10 mM Tris-HCl pH = 7.5. A test library amplification determined the number of PCR cycles necessary to meet Capture input requirements (200-500 ng per sample) using 5% or less of the prepped library, with test PCR reactions run on an agarose gel and yields quantified using image quantification software Image Studio Lite (LI-COR Biosciences). The Micro-C replicates in this manuscript were amplified with 8-10 PCR cycles for final library amplification using the KAPA HiFi HotStart ReadyMix (Roche #07958927001). Following library amplification, amplified libraries were purified to remove adaptor dimers, primers, and contaminants using AMPure XP beads (Beckman Coulter #A63880). Purified libraries were quantified via Fragment Analyzer and qPCR at the MIT BioMicro Center to determine library concentrations for pooling prior to Capture.

*Capture probe design (erythroid samples only)*
The probe panel was synthesized and purchased as a Custom Target Enrichment Panel from Twist Bioscience. A breakdown of the Custom Target Enrichment Panel is provided below. Only the BCL11A panel was used in this manuscript (**Fig. 1D**).

| Probeset Name | Twist Design ID | Region | Chromosome | Coordinates (hg38) | Panel Size (Mb) |
|---|---|---|---|---|---|
| Blood Panel 1 | TE-91601360 | ***BCL11A*** | **2** | **59,964,475-60,946,280** | **0.982** + 1.756 + 1.824 + 1.752 + 0.654 + 0.463 = 7.431 |
| | | *BACH2* | 6 | 89,233,475-90,989,896 | |
| | | *HBS1L-MYB* | 6 | 134,131,179-135,954,920 | |
| | | *HBB* | 11 | 4,457,550-6,209,242 | |
| | | *ZBTB7A* | 19 | 3,730,974-4,385,440 | |
| | | *KLF1* | 19 | 12,654,557-13,117,066 | |

*Capture of target loci*
Capture was performed in accordance with Twist Bioscience's Standard Hybridization Target Enrichment Protocol. Sample libraries were pooled in a 1:1 molar ratio across conditions, after which they were dried and mixed with Hybridization Mix (Twist Bioscience #104178), Custom Panels (Twist Bioscience #101001), Universal Blockers (Twist Bioscience #100578), and Blocker Solution (Twist Bioscience #100578). The library pool was hybridized to the biotinylated probe panel overnight, after which streptavidin beads (Twist Bioscience #100983) were used to pull down probes with hybridized ligated fragments and then washed (Twist Bioscience #104178) to remove unbound fragments. Another round of PCR amplified the target-enriched library using the Equinox Library Amplification Mix (Twist Bioscience #104178), with a test PCR included (as described above) to identify the number of amplification cycles necessary to meet sequencing requirements. With 3 μg of pooled input library for Capture, the RCMC samples generated in this manuscript needed 6 cycles of post-Capture PCR amplification. Following PCR

amplification, the Captured library was purified (Twist Bioscience #100983) and then quantified via both Fragment Analyzer and qPCR at the MIT BioMicro Center in preparation for sequencing submission.

##### Sequencing

Following qPCR quantification, post-Capture libraries across replicates were pooled in a 1:1 molar ratio. Pooled libraries were paired-end sequenced using two sequencing runs, one at 2x150 and one at 2x50 bp cycles, on a NovaSeq X system by the Broad Institute of MIT and Harvard's Walk-Up Sequencing services. Basecalls for NovaSeq output were performed using bcl2fastq v2.20.0.422.

##### Western Blot

Cells were harvested and lysed in RIPA buffer. Denatured protein samples were resolved on 4-15% Mini-PROTEAN® TGX™ precast gels (Bio-Rad) and transferred to PVDF membranes using the Trans-Blot Turbo system (Bio-Rad). Membranes were blocked in 3% BSA in PBST (PBS with 0.1% Tween-20) for 30 min at room temperature, and incubated overnight at 4 °C with primary antibodies against NFE2 (Abcam), STAG1 (Proteintech), STAG2 (Proteintech), or GAPDH (Proteintech). After washing with PBST, membranes were incubated with HRP-conjugated secondary antibodies and developed using ECL substrate (Bio-Rad). Signals were visualized using a ChemiDoc imaging system (Bio-Rad).

##### RNA sequencing

RNA-Seq for the NFE2 and NIPBL perturbation experiments (**Figs. 4-5**) was performed as follows. RNA extraction was performed from 0.5-1 million cells reserved from the same samples used for Micro-C using the Norgen Total RNA Purification Plus Kit plus gDNA removal column. Strand-specific RNA-Seq sequencing libraries were prepared by Novogene by poly(A) mRNA enrichment. Sequencing was performed on the NovaSeq X Plus platform, generating 150-bp paired-end reads.

#### Data Analysis

All processing and analytical code necessary to generate the analyses and results in this study from raw data is provided on GitHub (https://github.com/ahansenlab/erythroid_3D_genome).

##### Micro-C Processing

Paired-end reads were aligned with bwa-mem2 (v2.2.1) (4) with the -SP flag to the UCSC hg38 analysisSet genome and processed with pairtools (v1.1.2) parse2 (flags: --add-columns mapq --expand --report-position outer --min-mapq 30 --max-insert-size 150) (5). Reads from the same replicate were merged then deduplicated with pairtools dedup (flags: --max-mismatch 1).

Two sequencing runs were performed, the first at 50bp paired-end reads and the second at 150bp. Both runs were processed using the same pipeline. Replicate-wise .pairs files were subsequently merged between sequencing runs (pairtools merge) and deduplicated once more (pairtools dedup). Due to sequencing read mismatch between phases, Phase 1, Rep 2 was downsampled using pairtools sample 0.65 to yield comparable read depth to the second replicate. Read balancing was important in this case given that the Micro-C replicates represent different donors who may have intrinsic biological variability, and therefore no phase should have a read bias towards either replicate.

Final .pairs files were indexed with pairix (v0.3.8), and made into a .cool file with cooler (v0.10.3) cload pairs at 50bp bin size (6). Finally, .cool files were converted to .mcool files using cooler zoomify with the --balance flag up to 10Mb bin sizes.

##### RNA-Seq data processing

RNA-seq was aligned using the STAR aligner (v2.7.11b) (7) and deduplicated using samtools markdup (v1.0.1) (8,9). Count matrices were generated using subread featureCounts (v2.0.6) (10). Differential expression was performed using DESeq2 (v1.44.0) (11). Fold changes were shrunken using the 'ashr' method (12). Differentially expressed genes were considered to be those with adjusted p–value < 0.05 and shrunken log2(fold change) <-0.5 or >0.5. For NFE2 perturbation, the two biological donors for each condition (NFE2-KO and AAVS1-targeting) were considered replicates; for the NIPBL perturbation, the two guide conditions were considered replicates.

##### Micro-C reproducibility and read count reporting

Micro-C reproducibility in **Fig. S1C** was computed across all phases and replicates using HiCRep (v0.1.0) at 5kb binsize and 2Mb maximum genomic separation (13). Uniquely mapped read counts were obtained directly from pairtools stats after deduplication for each replicate. All read counts including mapped fraction, duplicated fraction, cis reads, and cis reads >1kb for all Micro-C datasets generated in this study are reported in **Table S1.**

##### Epigenomics processing and annotation

Sources for epigenomic datasets used to construct functional annotations in this study can be found in **Table S2**. All datasets were re-aligned to the hg38 analysisSet genome with bowtie2 (v2.5.4) (14) with flags --no-mixed --no-discordant for paired-end reads. Uniquely mapped reads were retained, and duplicate reads were removed with samtools. ChIP-Seq peak calling was performed using MACS2 (--BAMPE for paired-end ChIP and –BAM for single-end; narrowpeak mode for H3K27ac and CTCF and broadpeak mode for H3K4me1) (15). Cutoffs were determined using MACS2 –cutoff-analysis and verified by inspection. CUT&RUN peak calling was performed using SEACR (v1.1.3). bigWigs used for visualization and peak coverage analyses were made using deepTools (v3.5.6) bamCoverage (16).

Enhancers were denoted per-phase as the intersection of H3K27ac and H3K4me1 peak calls, except for P3 where H3K4me1 data was not available. In this case, H3K27ac alone was used to define enhancers. Promoters were denoted using the UCSC RefGene hg38 TSSs with a pad of 2kb upstream and 1kb downstream. CTCF was denoted per-phase as the intersection of called peaks with CTCF motifs in the hg38 genome. Motifs were generated using FIMO (17) over a Markov background with flags: –thresh 1e-3. To create unified enhancer and CTCF annotations across phases, enhancer and CTCF peak calls were merged across phases using bedtools (v2.27.1) merge (18). For enhancers, a pad of 500bp was used to join adjacent peaks.

ATAC-Seq peaks were from Martin-Rufino and Caulier *et al.*, representing 4 timepoints throughout erythroid differentiation using a similar three-phase culture system, which spans the P1, P2, and P3 phases in this study (19). ATAC-Seq and RNA-Seq from Ludwig and Lareau *et al.*, which analyze 8 timepoints through erythroid differentiation from bone marrow CD34$^+$ HSPCs, were used for visualization and for the linear regression in **Fig. 3B** (1). Briefly, ATAC-Seq and RNA-Seq from P123 were pooled for P1, P345 was pooled for P2, and P5678 was pooled for P3, and converted from hg19 to hg38 using UCSC LiftOver (20). Prior work has confirmed the similarity of these datasets to the P1-P3 cell types generated in this study (19).

#### Loop calling, classification, and quantification

Loops were called using Mustache (v1.3.3) at 1kb (flags: -r 1kb -pt 0.2 -st 0.92), 2kb (flags: -r 2kb -pt 0.2 -d 4000000), and 5kb (flags: -r 5kb -pt 0.2 -st 0.88 -d 4000000) binsizes (21). All loop calls were post-filtered at an FDR <0.01 to improve true-positive detection. Subsequently, all loop anchors were extended to 5kb, and loop calls were deduplicated with the priority 1kb > 2kb > 5kb, with a maximum of 2.5kb distance allowed between anchors that were called as duplicates.

Loops were classified according to the epigenomic annotations detailed above. Unless noted otherwise, loop classifications are inclusive. By this definition, loop anchors with multiple annotations were retained, and a single annotation was selected with the hierarchy promoter > enhancer > CTCF. That is, loop anchor overlapping a promoter with an enhancer mark is denoted as a promoter, an enhancer with CTCF is denoted as an enhancer, etc. In cases where loop classifications are labeled as exclusive, only loop anchors with a single classification are retained (all calls with multiple annotations are dropped). Loops were overlapped with annotation intervals using R's GenomicInteractions (v1.38.0) and InteractionSet (v1.32.0) (22,23).

Loops were quantified using AbLE (24) with the parameters gaussian_blur_pix=3, outlier_removal_radius_pix=3, and na_dist_to_center=1. Local backgrounds were computed at 100kb and quantification was performed at 10kb windows at 2kb binsize.

#### EryTF CRE and associated definitions

The eryTF CRE analyses in this study draw from the scATAC-Seq and TF-knockout experiments performed in Martin-Rufino and Caulier *et al.* (19). ATAC-Seq peaks of transcription-factor sensitive (TF) and erythroid-associated (ery) peaks were obtained from the Perturb-Multiome dataset (19). Subsequently, these were integrated with enhancer, promoter, and/or CTCF annotations as described above to generate the eryTF CREs analyzed in this study. Specifically, phase-unified enhancers without CTCF (enhancer annotation not overlapping promoters and CTCF) were intersected with eryTF peaks using bedtools. Similarly, ATAC-Seq peaks and accessible promoters (all not overlapping CTCF) were used as controls.

For the perturbation analyses (NFE2 and NIPBL), the classifications for candidate matchmakers were modified as follows. For NFE2, the subset of TF-sensitive elements that were only NFE2-sensitive were intersected with enhancers not overlapping CTCF. For STAG2 and NIPBL analyses, the set of all enhancers not overlapping CTCF was used (i.e. not necessarily eryTF-annotated). This allowed more direct comparison with the STAG2 U937 and G1E annotations, which lack functional annotations such as the eryTF annotations. Clustering and matchmaker annotation for erythroid perturbation datasets were computed on the differentiation Micro-C due to the fact that the perturbation Micro-C was 0.3-0.5x the read depth of the differentiation Micro-C, which may affect the rank order or clustering of loci. Thus, maintaining a consistent clustering and matchmaker annotation ensures that the same loci are represented consistently across datasets.

U937 enhancer annotations were determined using H3K27ac ChIP-Seq and promoter annotations were determined using H3K4me3 ChIP-Seq (25). CTCF annotations were from the erythroid datasets analyzed in this study. G1E enhancer and promoter annotations were from Zhang *et al.*, Nature 2019 (26). G1E CTCF annotations were from (27). In both U937 and G1E analyses, the enhancer annotations as described above were filtered to exclude overlaps with promoter and CTCF annotations. The anti-insulation score, k-means clustering, and matchmaker annotation were then computed on these filtered enhancers exactly as described for the erythroid differentiation Micro-C (see below), except that G1E Hi-C matrices were binned at 5kb.

#### Matchmaker analyses

Anti-insulation score was computed as follows. For each genomic element to be scored, the observed-over-expected matrix 100kb on either side of the input genomic element (hence, "input element") at 2kb binsize was obtained using cooltools (v0.7.1) ObsExpSnipper, excluding the first two diagonals as in the cooltools default (28). Next, the 3D genomic contact intensity in the "crossing" quadrant and the "same-side" triangles (see **Fig. 2G**) were each summed and the anti-insulation score was computed as the sum of the crossing quadrant divided by the sum of the two same-side triangles. Majority-NaN regions (>80% of pixels) were dropped.

To classify loops, all loops within the matrix bounds were retrieved. Loops were classified as same-side if anchors were on either both on the left or both on the right of the input element (in genomic coordinates). Loops were classified as crosser loops if one anchor was on the left and one anchor was on the right of the input element. Observed/expected intensities at loops were computed by taking a 5kb window around each loop anchor's middle point (i.e. a 10kb x 10kb square). In the erythroid perturbation Micro-C, loop calls from the differentiation Micro-C were applied to the loop score quantification in the perturbation Micro-C. Loop analyses were not performed in the other cell types (G1E, U937).

After computing all anti-insulation scores, all input elements were deduplicated within 10kb by selecting the element with the maximum anti-insulation score. This was performed in order to avoid creating artificial biases in the distribution from clustered elements. Since eryTF CREs are defined from ATAC peaks, this avoids over-representing single enhancers with multiple accessible sites. All matchmaker analyses excluding the ridge regression use this deduplicated set.

All local 3D genome pileups were performed using coolpuppy (29) at 5kb binsize.

#### Insulation, boundaries, compartments

All insulation and boundary analyses were performed using cooltools insulation. Fine-scale insulation analysis (**Fig. 1C** and **S2D**) was performed at 2kb with 20kb window sizes. To define high-confidence CTCF TADs, cooltools insulation was run at 10kb binsize with 100kb windows and TADs were identified as described in the cooltools documentation. TADs overlapping CTCF were identified with bedtools pairToBed with the flag -either. This TAD boundary definition aligns with a similar analysis done on fountains (30), where the position of fountains relative to TAD boundaries was computed.

For coarse-scale (100kb and 10kb) compartment analyses in **Figs. 1** and **S2**, 1st eigenvectors were used to define A/B compartments using cooltools. For eigenvector inputs into the regression in **Fig. 3B**, 2kb-binsize Micro-C eigenvectors were computed using POSSUMM (31).

##### Motif searches

Motif searches were performed using HOMER (v5.1) findMotifsGenome with -size 100 (32). The test sets (gained and lost boundaries) were strong boundaries (annotated by cooltools insulation at 2kb binsize/20kb window) that changed annotation between the respective phases, subsequently intersected with ATAC peaks. The background set was all boundaries pooled across phases, intersected with ATAC peaks.

##### Gene expression analyses

For erythroid differentiation gene expression analyses, the scRNA-Seq of the control cells (non-targeting gRNA or no treatment) from Martin-Rufino and Caulier *et al*. (19) were re-analyzed. Seurat AggregateFunction() was used to get pseudo-bulked populations for P1, P2, P3, and eosinophil/basophil/mast cell precursors as an off-lineage control. P3 marker genes were extracted using Seurat (v5.4.0) FindMarkers() against the other populations (33). Cutoffs of $p<0.01$, 10% expressed in the cell type of interest, and $\log2FC>0.25$ were used to filter marker genes. For pseudo-bulked expression, log-normalized transcription>0.1 was considered expressed. Further, for the comparison with P3-comparable gene expression and P3 marker genes in **Fig. S13C**, the comparable gene expression set was the set of P3-expressed genes with expression higher than the $25^{th}$ percentile of P3 marker genes, which was chosen to match median expression levels between the two sets.

All statistical testing of distributions was performed using the Kruskal-Wallis test with post-hoc Mann-Whitney U tests (p-values corrected by the Bonferroni correction) if the comparison was multiway, or a Mann-Whitney U test if the comparison was two-group. Enrichment analyses were computed with odds ratio testing using Fisher's exact test (see below).

Analyses linking marker or expressed genes to matchmaker loops (**Fig. 4D-E**) or linking gene expression to erythroid loops (**Figs. 2C** and **S3F-G**) were performed as follows. Promoter-associated chromatin loops were assigned by first removing redundant promoter annotations arising from duplicate transcripts while retaining distinct transcription start sites. Then, loops were assigned to promoters by anchor, if either anchor overlapped the TSS window (2kb upstream and 1kb downstream). Promoter–loop assignments were merged with precomputed AbLE score measurements to associate each promoter-linked loop with its corresponding loop strength. For analyses of gene expression as related to loops only (**Figs. 2C and S3F-G**), if multiple loops overlapped, only the loop with the best anchor overlap was selected for each promoter and the top and bottom 5% of gene expression values were excluded.

##### Odds ratio testing

The odds ratio tests in **Fig. 2A** (odds of observation of erythroid TF genomic elements on loop anchors) were performed by computing the overlap of eryTF peaks and loop anchors using bedtools intersect. Similarly, the background set was a random sample of ATAC peaks overlapped with loop anchors. Thus, the odds ratio tests the likelihood of observing eryTF peaks on loop anchors vs. ATAC peaks on loop anchors. Statistical significance represents corrected false discovery rate (Benjamini-Hochberg) for multiple comparisons.

Odds ratio tests in **Figs. 4B, S13C, 5I** and **S15E** were all performed to test the enrichment of gene proximity to eryTF CRE clusters. Genes were compared across matchmaker clusters by finding the closest eryTF CRE within 100kb and reporting its cluster, if at least one was detected. Thus, the gene observation odds ratio tests ask: is the proportion of the test set that is near each eryTF CRE cluster significantly higher than the proportion of the background set near each eryTF CRE cluster?

For all human erythroid analyses, the background was the set of accessible promoters, upon verification that the accessible promoters were a near-superset of the genes being analyzed. This aimed to control for active gene density between matchmaker clusters. For the G1E analysis, the background was the set of promoter annotations under the assumption that the promoter annotations were a superset of the differentially expressed genes. Statistical testing was performed using Fisher's exact test.

#### Aggregate peak analyses

All 1D aggregate peak analyses were performed using deepTools in reference point mode and report the mean value within every genomic bin (flag –averageTypeBins mean). The exception is the analysis of anti-insulation across H3K4me1 and H3K27ac peak strengths in **Fig. S8B-C**, where scale-regions mode at 10bp binsize was used to compute the mean signal across the entire peak for all peak calls. Then, peak calls with a higher signal than the 10th percentile of the signal of enhancers that overlapped eryTF CREs were used to define boundaries for the quantiles shown in **Fig. S8B-C**, in order to represent peaks of similar strength to eryTF CREs. The four quantiles shown are even quantiles by peak occupancy signal.

#### Ridge regression

The Ridge regression (**Fig. 3B**) was used as an explanatory indicator of the features associated with matchmakers via anti-insulation score. Generally, the regression attempts to predict the anti-insulation score of a given eryTF CRE given its epigenomic context. First, deepTools was used to obtain matrices of 14 epigenomic/transcriptomic features across all three phases used as inputs (**Fig. S7**, **Table S2**), using computeMatrix in –reference-point mode centered at each of the eryTF CREs. The last three inputs were 2kb-binsize eigenvectors, which were computed using POSSUMM as described above. All epigenomic/transcriptomic inputs (i.e., all except the eigenvectors) first underwent a log10(X+1) transform. Then, all inputs were binned according to distance from the center (i.e. the distance from the eryTF CRE) in ten 10-kb bins. That is, the first bin represents all signal that is 0-10 kb on either side of the center, the second bin represents all signal that is 10-20 kb on either side, etc. Finally, the binned inputs were normalized using sklearn's StandardScaler. The regularization value (alpha) was selected by running 200 random train-test splits and constructing a distribution of alphas under which the MSE was minimized (Fig. S7A), although features were robust to variation of alpha within the range of this distribution (**Fig. S7C**). Then, the model was fit on the full dataset of anti-insulation score for each phase. Feature coefficients are plotted per bin to show the predictive values of each feature as a function of genomic distance from the eryTF CRE.

#### Polymer simulations

All polymer simulations were performed with polychrom (v0.1.0) (34). To assess the effect of targeted cohesin loading, the simulated locus from Jusuf *et al.* (24) was modified to incorporate targeted cohesin loading. The chromosome was modeled as a chain of 70,000 monomers, where a 2,000-monomer locus was repeated 35 times. This locus contained CTCF stall sites and CREs which exhibit affinity-mediated interaction. The configuration, stall probabilities, and CRE affinity were exactly replicated from Jusuf *et al.* (24). A 1D simulation of loop extrusion was used to obtain cohesin positions across the duration of the simulation, as described previously (24,35). Briefly, cohesins randomly load and dissociate from DNA with fixed probabilities. If one of cohesin's two motor subunits became stalled at a CTCF site, the other motor subunit of the cohesin was allowed to continue extruding until it was either stalled at another CTCF site or until the entire cohesin dissociated from the DNA.

Targeted loading was modeled as an increase in cohesin "birth" probability, i.e. an increased likelihood of cohesin loading, at the loading site monomers compared to the other monomers. Thus, the loading bias represents the fold likelihood of cohesin loading at a given monomer of the loading site as compared to the rest of the polymer. The total number of cohesins was kept constant. Three loading sites were simultaneously tested in each region (configuration described in **Fig. S11A**). To model chromatin compaction, the chromatin volume density parameter was modulated as described in Figs. 3F-G and S11C-D. The baseline chromatin volume density was 0.3 and increased compaction conditions were 0.45 and 0.6.

Loop quantifications for simulated maps were performed as follows. Polymer simulation contact maps were generated from the 35 repeats of the 2Mb region on the 70Mb chromosome, across all simulation blocks. For each simulation, contact maps were calculated across all blocks and normalized by both the number of blocks and the number of genomic regions sampled. Loops were defined from the positions of convergent CTCF sites, non-convergent CTCF site pairs, and CRE elements in Jusuf *et al.* (24).

Simulated contact maps were generated using polychrom's monomerResolutionContactMapSubchains function with a contact radius cutoff of 3 units. Estimations of loop strength and associated error were quantified using a leave-one-block-out jackknife procedure on each loop. For each simulation block, the corresponding contact map was excluded and the remaining blocks were used to generate a leave-one-out (LOO) contact map. For each loop, observed contact frequency was calculated as the mean signal within a 10kb x 10kb square window centered on the loop position. For each loop, jackknife estimates of observed contact frequency and O/E enrichment were collected across all leave-one-out iterations. Final loop strengths were reported as the mean of the jackknife estimates, and uncertainty was quantified using the jackknife standard error. In **Fig. S11D**, where maps at different compaction levels are compared directly, contact matrices were balanced using the Sinkhorn-Knopp algorithm to normalize for differences in total contacts, as more compacted DNA will exhibit higher contact probabilities on average.

#### Visualization

All contact maps with epigenomic tracks were visualized using a modified version of CoolBox (v0.3.8) (36). All other visualizations used matplotlib (v3.8.0), seaborn (v0.13.2) or R (v4.0.2) (base or ggplot2).

#### Data and Code Availability

Data generated for this study (Micro-C and RNA-Seq) can be found on GEO under accession numbers GSE341414, GSE341565, GSE341752 and GSE341753. Publicly available datasets re-analyzed for this study can be found in **Table S2**. Code for analyses and figure generation can be found on GitHub at https://github.com/ahansenlab/erythroid_3D_genome.

### Supplementary Figures

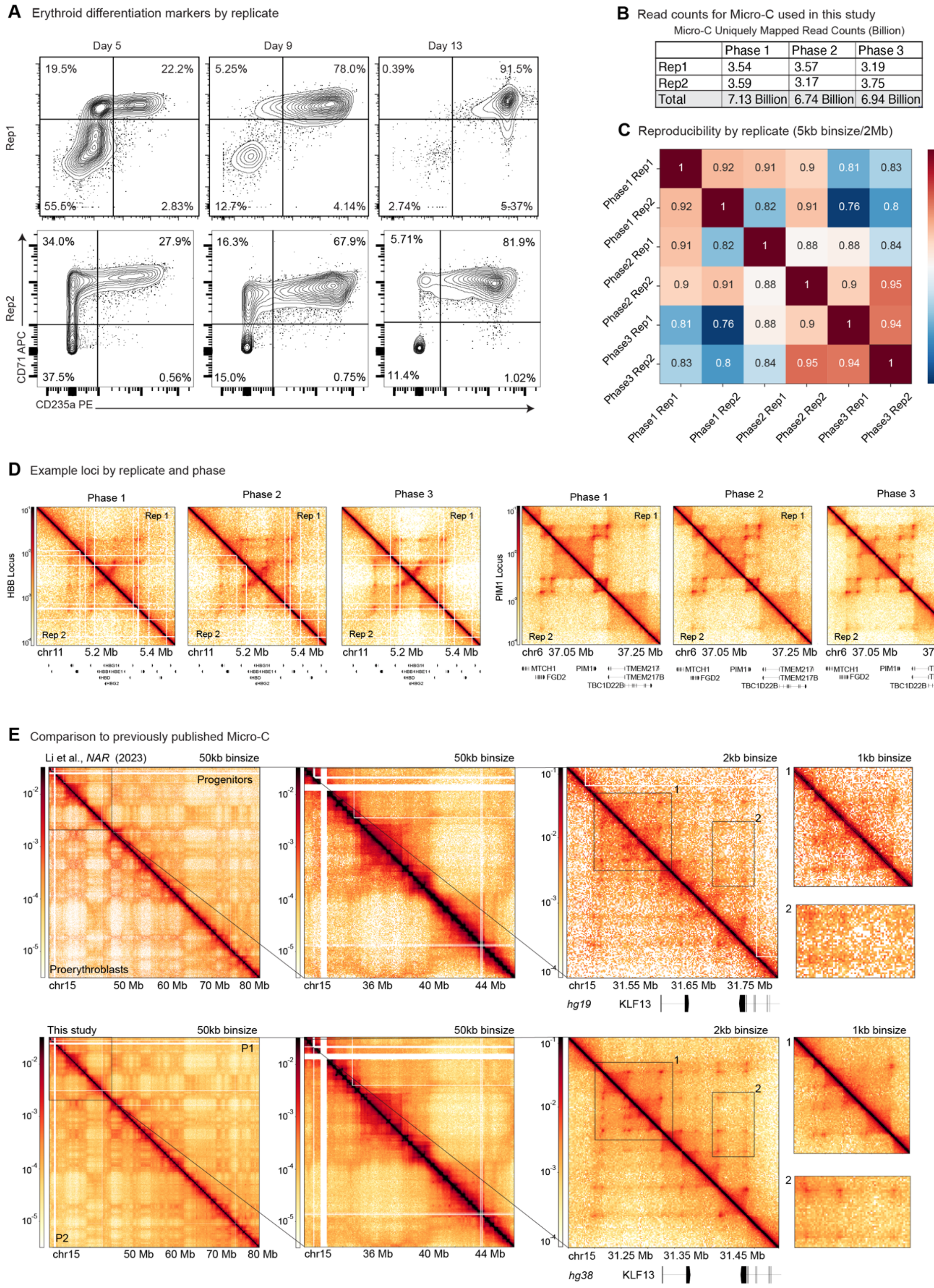

**Supplementary Figure 1: Overview of erythroid differentiation Micro-C**. (**A**) Differentiation state at each phase of erythroid differentiation assessed by fluorescence-activated cell sorting for CD235a and CD71. Replicates represent different donors. (**B**) Uniquely mapped read counts for each phase and replicate used in this study. Full statistics are reported in **Table S1**. (**C**) Micro-C reproducibility by replicate as determined by HiCRep at 5kb binsize up to 2 Mb genomic distance. (**D**) Examples of key erythroid loci in each replicate at each differentiation timepoint. Left: beta-globin locus (chr11:5007550-5509242) at 3.2kb binsize. Right: *PIM1* locus (chr6:36967226-37358383) at 2kb binsize. (**E**) Comparison of the *KLF13* locus in the deepest Micro-C across erythropoiesis to our knowledge (37) and our dataset. Coordinates were converted from hg38 (chr15:31150000-31550000) to hg19 (chr15:31442203-31842203) using UCSC LiftOver.

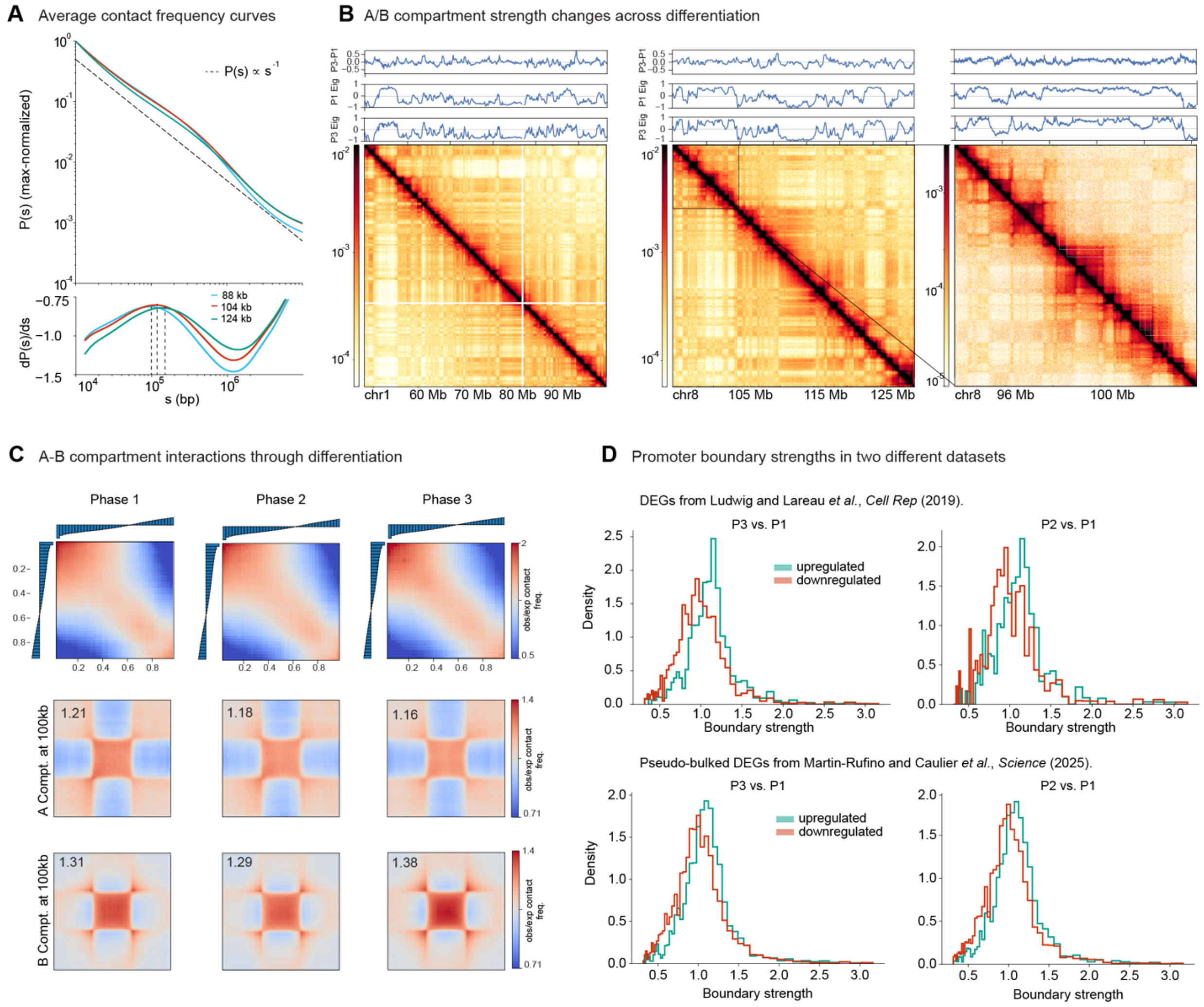

**Supplementary Figure 2: Compartment and boundary changes across erythropoiesis. (A)** Top: Contact probability as a function of genomic separation (*P(s)*) across differentiation timepoints, computed at 2kb binsize. Dashed line represents *P(s)* ∝ $s^{-1}$ scaling. Bottom: First derivative of the *P(s)* curve with respect to *s*. Dashed lines, with annotation in legend, represent the s value where the first derivative is maximized. **(B)** 1st eigenvectors of the Micro-C matrices, which typically represent A/B compartmentalization at two example loci: left, chr1:48000000-102000000 at 100kb binsize, right, chr8:93846768-104070949 at 100kb binsize with inset chr8:93846768-104070949 at 10kb binsize. P1/P3 eigenvectors, as computed by cooltools at the denoted binsizes, and the difference vector are shown. **(C)** (Top) saddle plots of A/B interaction strength. Top left represents the strongest BB interactions, and bottom right represents the strongest AA interactions. (Middle) Rescaled A-compartment pileup; (bottom) Rescaled B-compartment pileup. All analyses were performed at 100kb binsize and for compartment pileups, only compartments longer than 500kb were considered. **(D)** Boundary strengths, as annotated by cooltools insulation at 2kb binsize and 20kb window size, at upregulated and downregulated genes from two different studies.

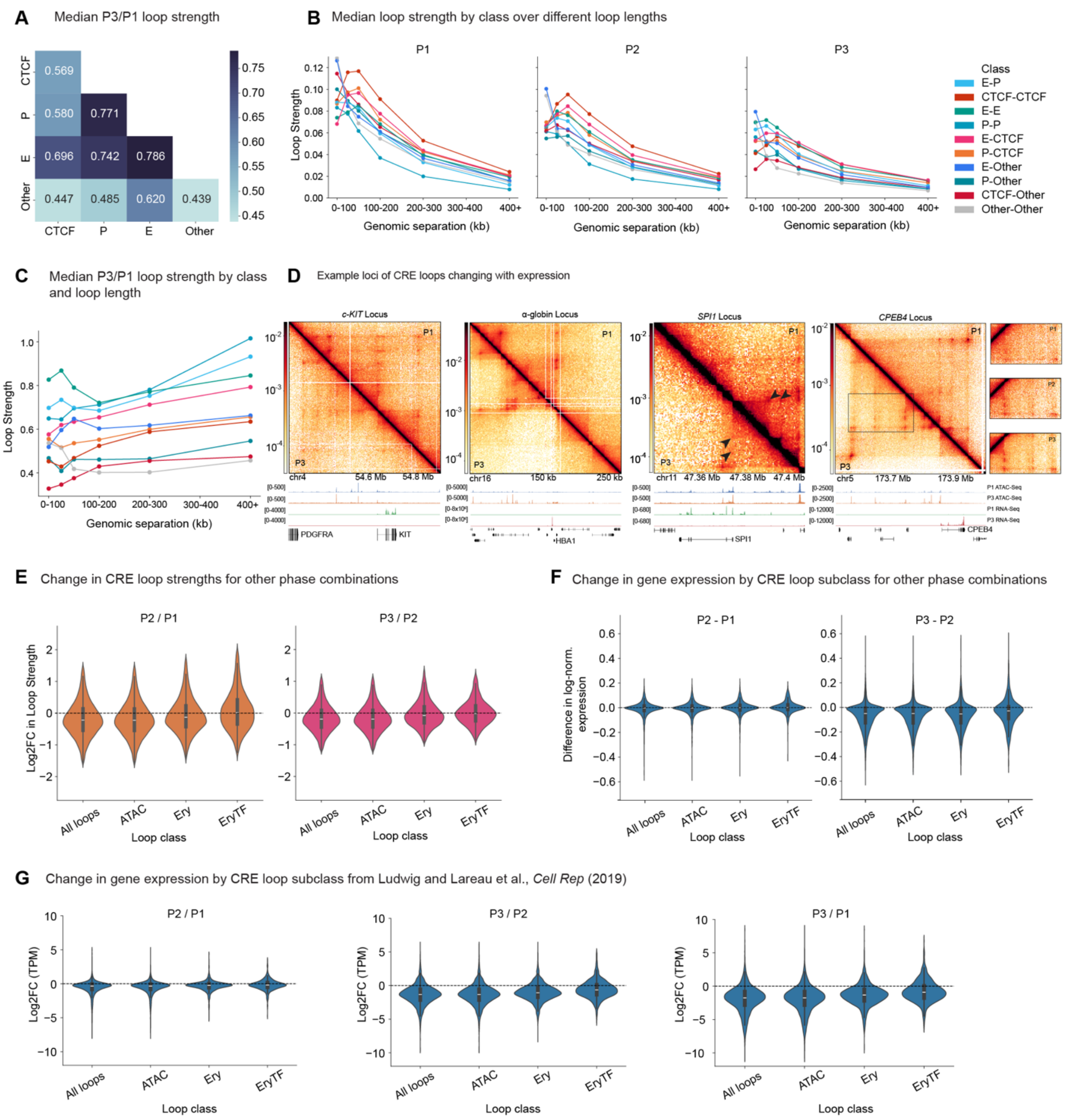

**Supplementary Figure 3: Loop strengths relate to gene expression changes during differentiation. (A)** Median change in loop strengths (AbLE) for each loop class (inclusive of multiple classes, see Methods) between P1 and P3. **(B)** Median loop strength at each phase by inclusive loop class, stratified by loop length (genomic separation between the loop anchors). Each point represents the median loop strength of loops in the indicated distance interval. **(C)** Median change in loop strength by class stratified by loop length. **(D)** Example loci of CRE loops that trend with gene expression. From left to right: *KIT* locus (chr4:54257267-54940783 at 2kb binsize), α-globin locus (chr16:50000-277522 at 1kb binsize), *SPI1/PU.1* locus (chr11:47343511-47410593) at 500bp binsize, *CPEB4* locus (chr5:173570169-174026097; inset chr5:173670000-173830000 and chr5:173600000-173900000) at 1kb binsize. Arrows/insets denote changing loops. **(E)** Loop strengths as in **Fig. 2C** (top), for other phase combinations. **(F)** Changes in transcription as in **Fig. 2C** (bottom), for other phase combinations. **(G)** Changes in transcription as in **Fig. 2C** (bottom), for a different expression dataset (as referenced in **Fig. S2D**).

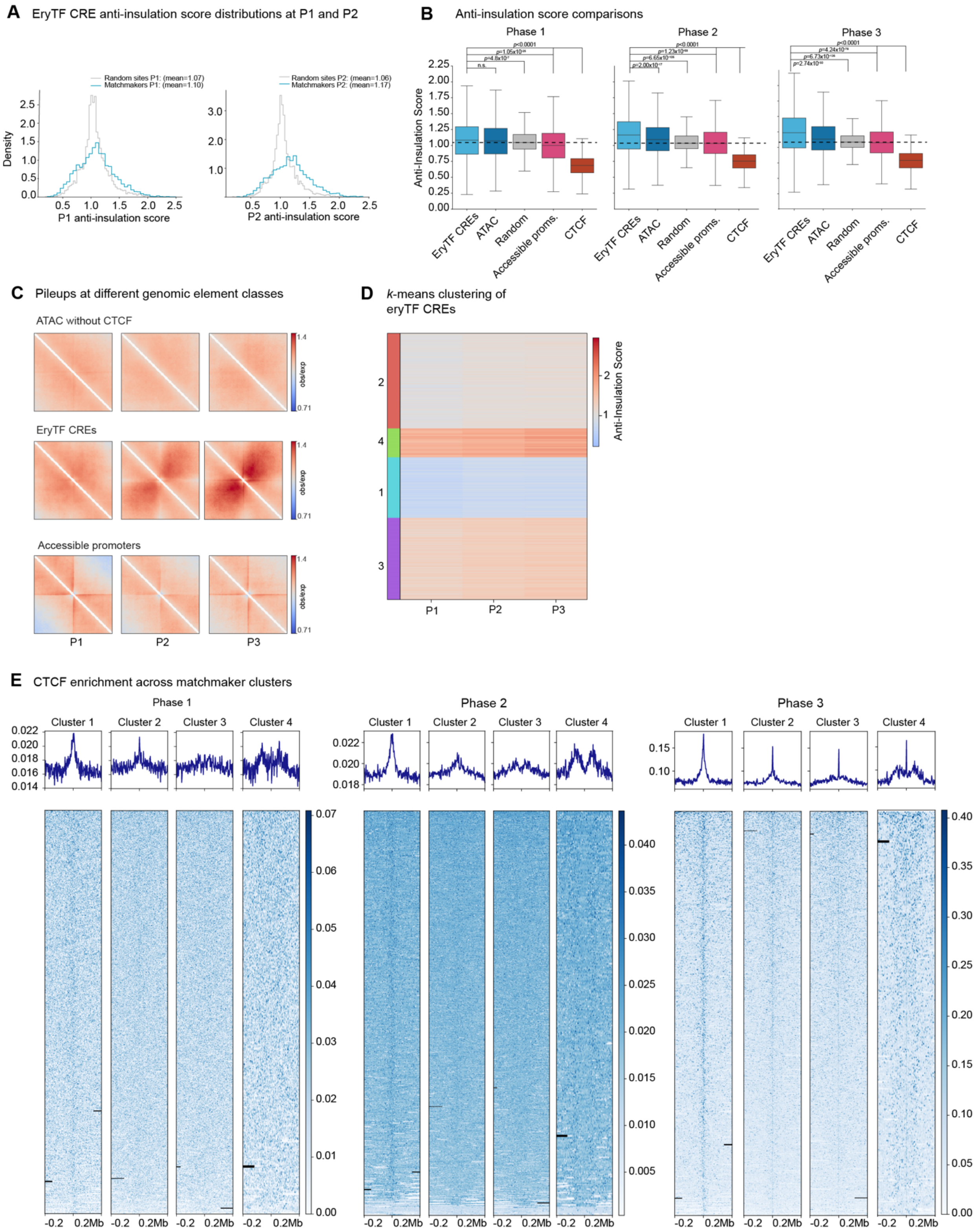

**Supplementary Figure 4: Characterization of anti-insulating eryTF CREs. (A)** As in **Fig. 2G**, anti-insulation scores for eryTF CREs in P1 and P2 Micro-C. Random sites were generated by shuffling ATAC intervals into the hg38 genome using bedtools shuffle. For P1, p=0.0002 and for P2, p=9.64x10$^{-60}$ for the difference in means by the Mann-Whitney U Test (two-sided). **(B)** Anti-insulation score distributions across phases for eryTF CREs (n=3324), ATAC peaks without CTCF (n=4687), random genomic sites (n=27670), accessible promoters (n=3513), and high-confidence CTCF boundaries (insulation boundaries at 10kb binsize/100kb window size with CTCF annotation) (n=3806). ATAC peaks without CTCF and accessible promoters were randomly sampled to 5000 elements prior to anti-insulation score calculation due to large sample sizes, and reported values indicate loci with valid anti-insulation scores (see Methods). All elements are deduplicated within 10 kb to avoid bias from clustered elements. Labeled p-values indicate the results of post-hoc two-sided Mann-Whitney U Tests adjusted for multiple comparisons by the Benjamini-Hochberg method. **(C)** Pileups at all three phases of ATAC peaks without CTCF, eryTF CREs, and accessible promoters, with 200kb window size at 2kb binsize. Each of these groups was randomly sampled to have a maximum of 5000 elements per group. **(D)** *k*-means clustering of eryTF CRE anti-insulation score, considering all phases. Clusters are annotated on the left. **(E)** Pileup of CTCF from each phase (see **Table S2**) at 2kb binsize and 200kb flank width centered at the eryTF CREs. Black lines indicate masked NaN values.

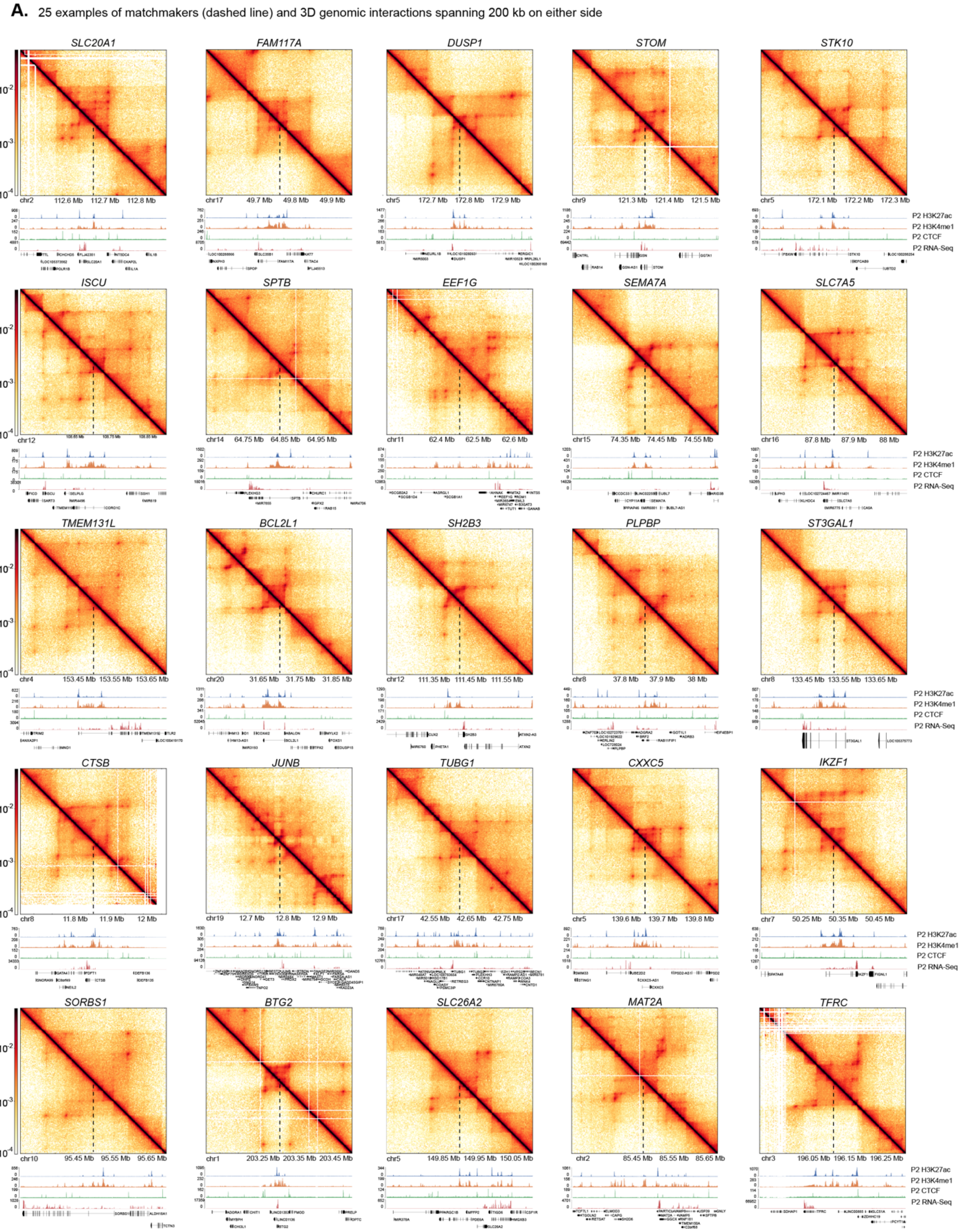

**Supplementary Figure 5: Examples of matchmakers and surrounding 3D genomic structure.** 25 example matchmakers identified via anti-insulation score. Top right is P1 and bottom left is P3. Dashed lines indicate the center of the locus, where the matchmaker was annotated. All visualizations are 200 kb on either side of the matchmaker. Titles indicate a representative gene name contained within the locus, selected by inspection based on expression level in P3 relative to other genes in the locus.

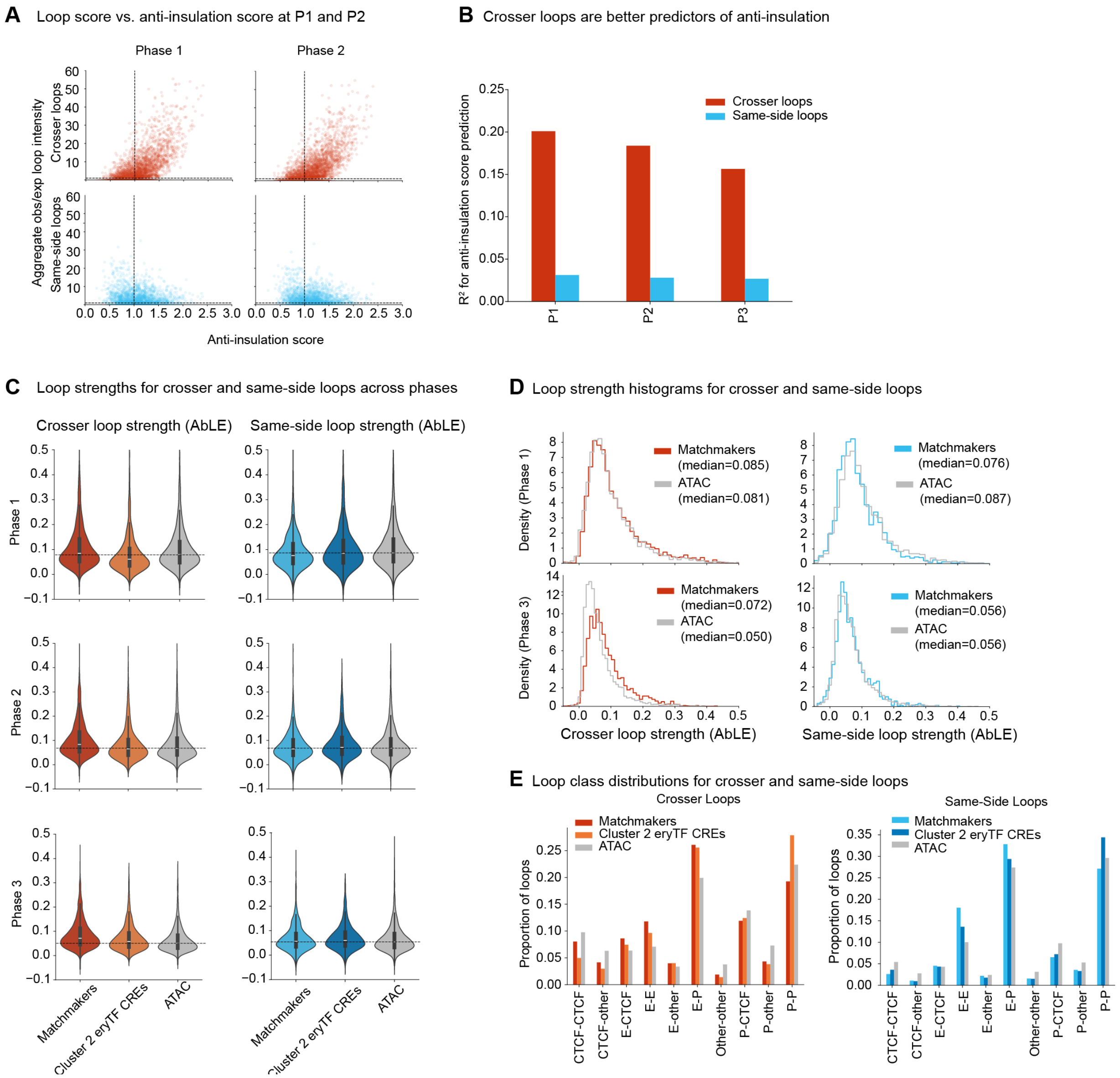

**Supplementary Figure 6: Matchmakers are characterized by strong crosser loops compared to same-side loops. (A)** Observed/expected contact at 10kb x 10kb windows centered around crosser and same-side loops as in **Fig. 3A**, but for P1 and P2 anti-insulation and loop scores. Loop scores are summed over all crosser or same-side loops within 100kb of each eryTF CRE. **(B)** $R^2$ coefficient of determination for crosser vs. same-side observed/expected intensity (summed per locus) against anti-insulation score. **(C)** Crosser and same-side loop strengths (via AbLE) for matchmakers (clusters 3-4 of eryTF CREs), cluster 2 of eryTF CREs, and ATAC-Seq peaks not overlapping CTCF (sampled to the size of eryTF CREs). **(D)** Histogram of crosser and same-side loop strength as in **(C)**, for matchmakers vs. ATAC peaks without CTCF at P1 (top) and P3 (bottom). (**E**) Inclusive loop class distributions (see Methods) for each of the three classes in (**C**). For matchmakers, n=1712 same-side, n=4110 crosser; for ATAC, n=3340 same-side, n=4023 crosser.

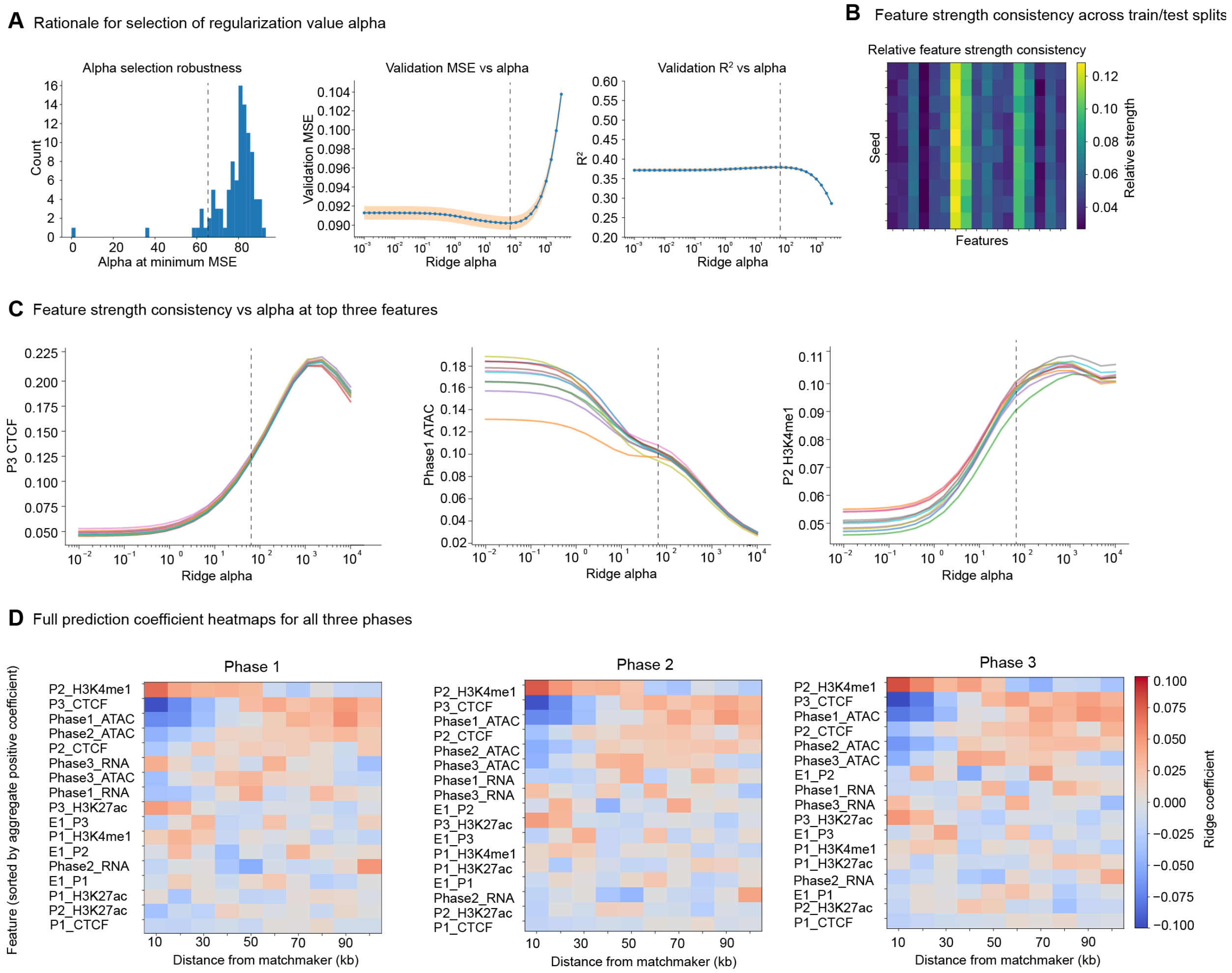

**Supplementary Figure 7: Parameter selection underlying ridge regression model. (A)** (Left) Regularization parameter alpha that minimizes the validation mean squared error (MSE) on 200 random 80:20 train-test splits of the data, across a range of 100 alpha values logarithmically spaced from $10^{-4}$-$10^{3}$. (Right) MSE and $R^2$ for prediction of anti-insulation score, across a range of 40 alpha values logarithmically spaced from $10^{-2}$-$10^{3.5}$. Shaded area indicates standard error on the mean across 50 random seeds for 80:20 train-test splits. The dashed line indicates the selected alpha. **(B)** Consistency of feature strengths across 10 different random data subsets. **(C)** Feature strengths of the top three features across a range of alpha values. Lines represent 10 random train-test splits to indicate feature stability as a function of alpha. The dashed line indicates the selected alpha. **(D)** Full coefficient heatmaps across distance bins, as in **Fig. 3B**, for all three phases with complete feature name labels.

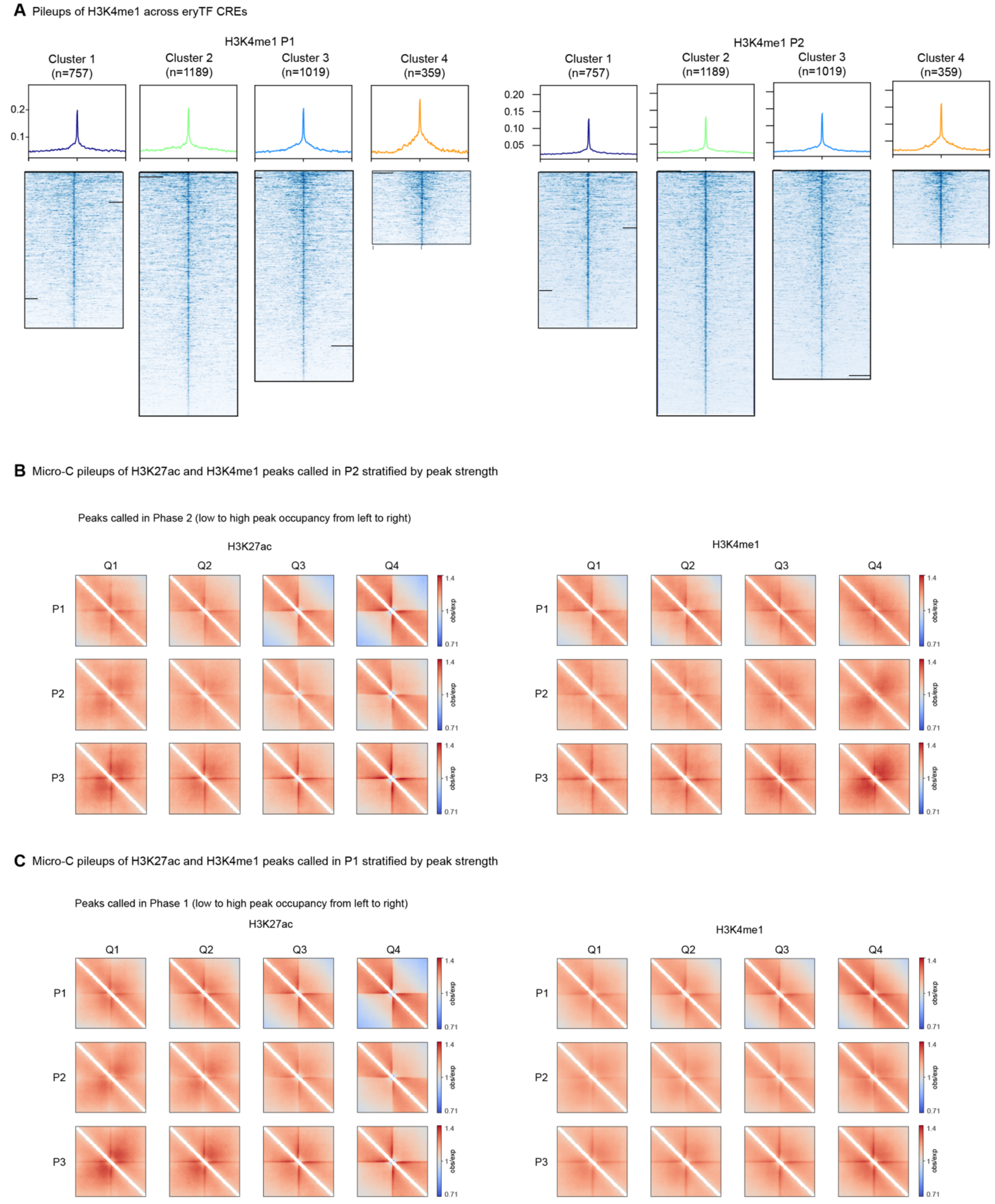

**Supplementary Figure 8: Matchmakers are characterized by strong H3K4me1. (A)** Aggregate peak analyses of H3K4me1 called in P1 and P2 across the 4 clusters of eryTF CREs. Black lines indicate masked NaN values. **(B-C)** Pileups centered at peaks called in H3K4me1 (in P2 (**B**) and P1 (**C**)) and H3K27ac without any other filtering (e.g. CTCF removal, promoter removal, etc.) evenly split across the range of 10th to 100th percentile of mean peak signals found at eryTF CREs (see Methods). Top to bottom: P1 to P3. Left to right: Low to high peak signal. Mean peak signal was determined using deepTools across the length of the peak call as annotated by MACS2 (see Methods for peak calling description).

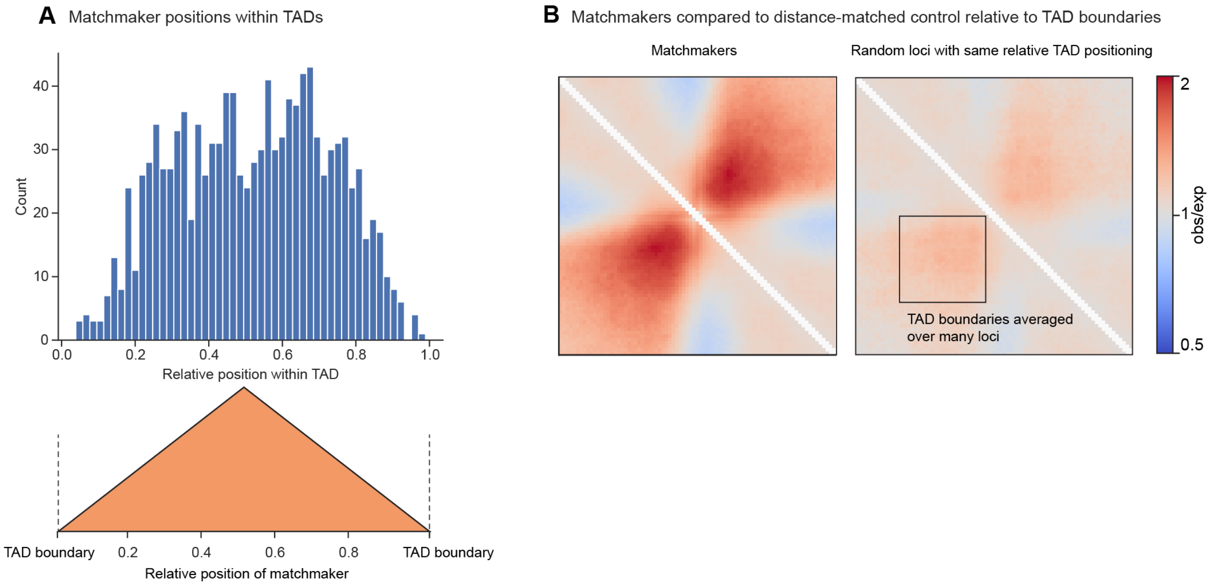

**Supplementary Figure 9: Matchmakers are not artifacts of topologically associated domains (TADs). (A)** Position distribution of matchmakers within TADs annotated by cooltools insulation as boundaries called with binsize 10kb and window 100kb containing a CTCF site (ChIP peak/motif) on at least one side. **(B)** Pileup of matchmakers as compared to 1000 random points position-matched relative to TADs using the distribution from (**A**), both plotted in P3 Micro-C.

**A.** 25 random examples of points within TADs and 3D genomic interactions spanning 200 kb on either side

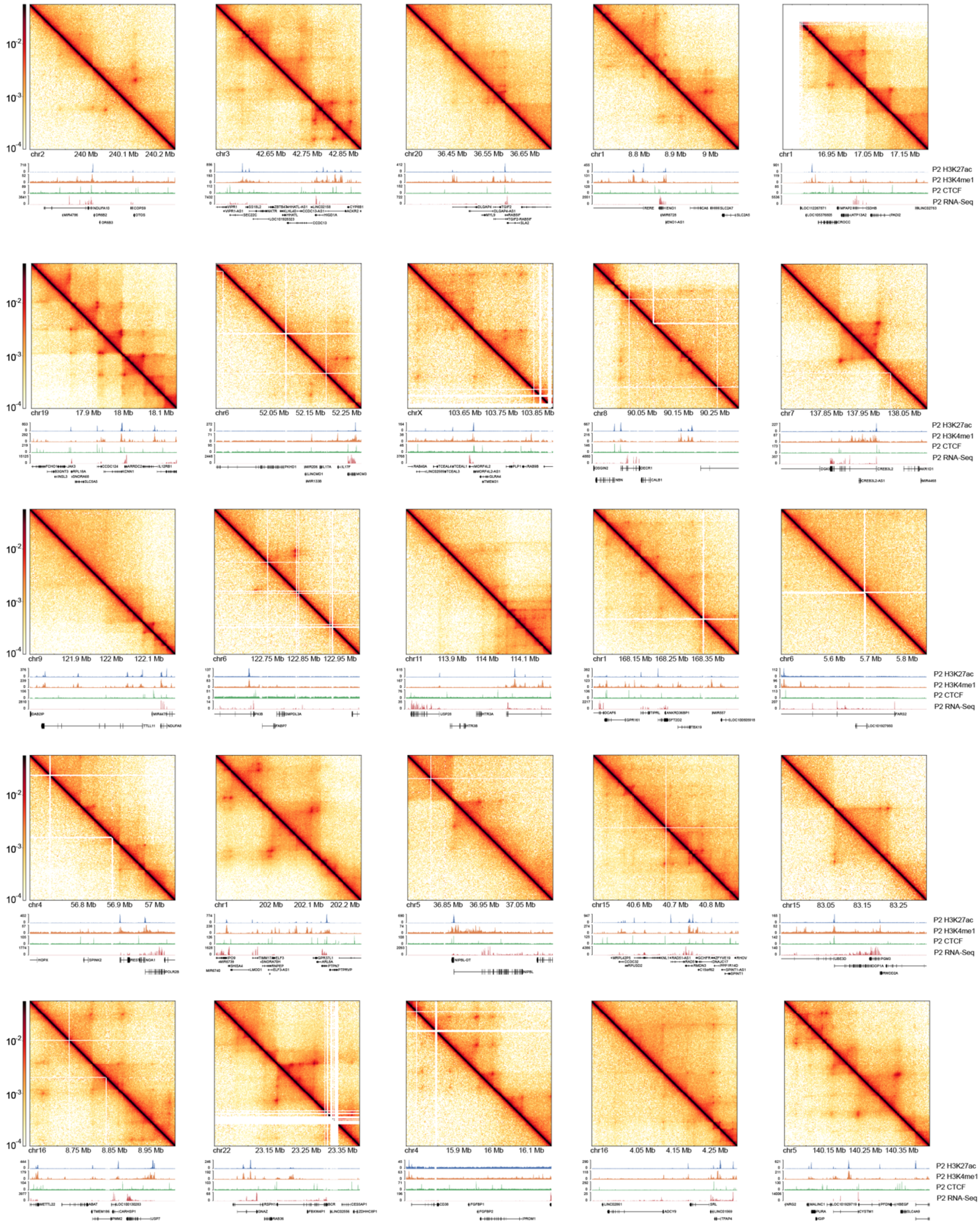

**Supplementary Figure 10: Examples of random positions distance-matched to matchmakers within TADs.** 25 random examples selected by applying the distribution of matchmaker positions within TADs to all TADs as in **Fig. S9B**. Top right is P1 and bottom left is P3.

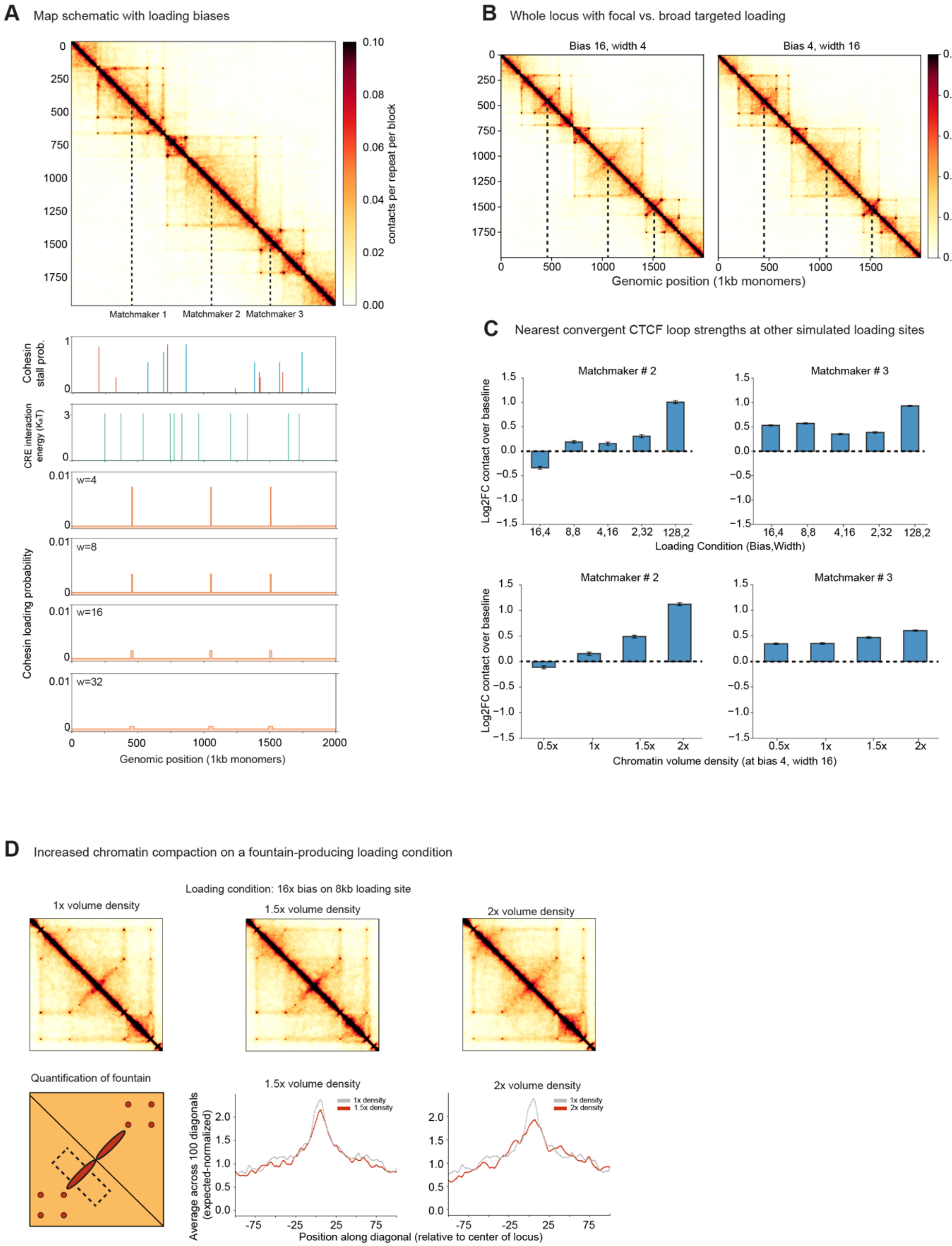

**Supplementary Figure 11: Polymer simulation overview and further characterization of preferential loading at matchmakers.** **(A)** Modification of the simulated locus from Jusuf *et al*. (24) to incorporate preferential cohesin loading at three sites in the 2Mb region. The depicted map is at baseline, with no loading bias. Top to bottom tracks show CTCF (left-pointing sites in blue and right-pointing sites in red), CREs, and examples of matchmaker-preferential loading regions. In this schematic, the number of SMC complexes that preferentially load on each matchmaker remains constant, but the width of the matchmaker changes, yielding different effective loading biases. **(B)** Simulated locus map with focal (left) and broad (right) loading. **(C)** Change in observed loop strengths at the two loading sites not shown in **Fig. 3F-G** under conditions of increased loading (top) and increased compaction (bottom). **(D)** (Top row) A condition that shows a discernible fountain at baseline chromatin volume density (16x cohesin loading bias over 8kb) under 1.5x and 2x volume density. (Bottom row) intensity profiles across the fountain in both conditions. Maps were balanced and O/E normalized for equal comparison.

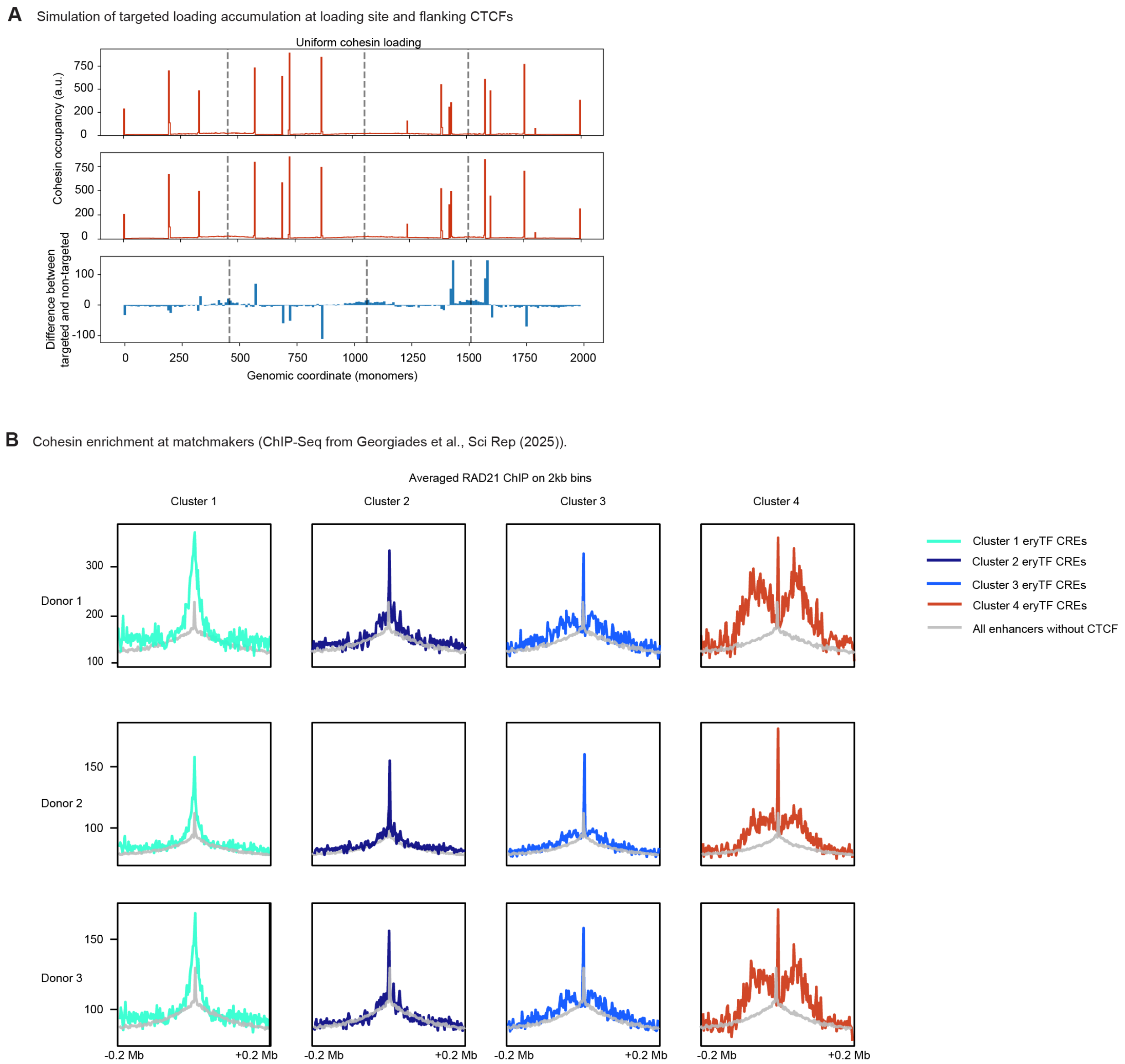

**Supplementary Figure 12: Cohesin accumulates at matchmaker-flanking sites. (A)** 1D simulation of cohesin under non-targeted and targeted cohesin loading (16x loading bias per loading monomer relative to random monomers, on 8-monomer loading pad) at the same locus as in Fig. 3 and S11. Bottom shows the difference between targeted and non-targeted. **(B)** RAD21 ChIP-Seq from human donor erythroid cells (38) averaged over each eryTF CRE cluster (2kb binsize and 200kb flank). Rows represent one replicate from three different biological donors.

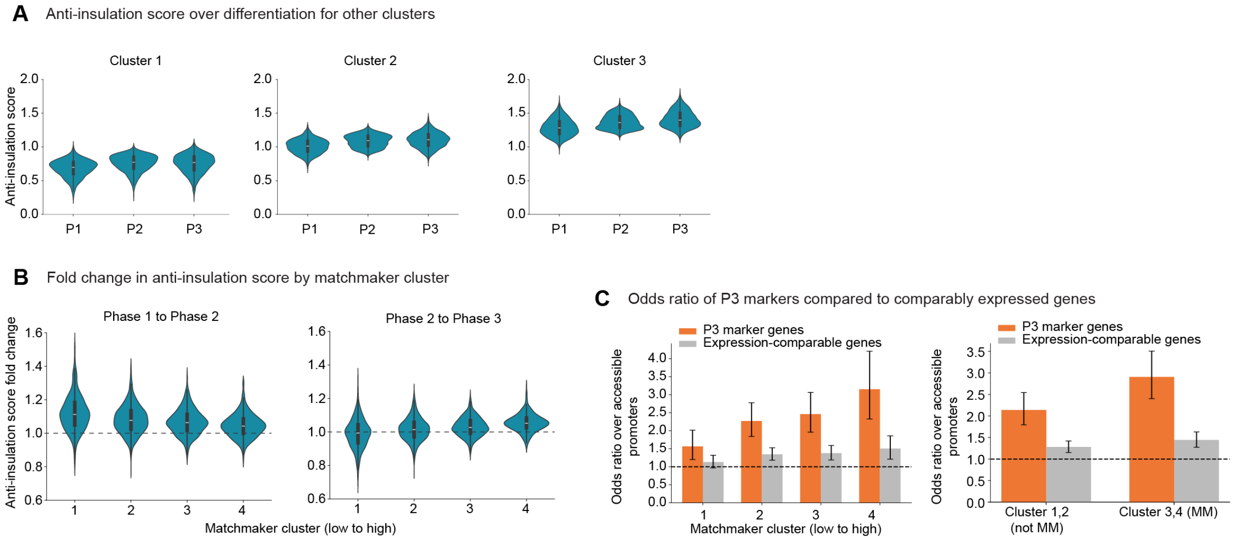

**Supplementary Figure 13: Matchmakers increase in anti-insulation over differentiation and are related to erythroid gene expression.** (**A**) Anti-insulation score distributions by phase as in **Fig. 4A**, for clusters 1-3. (**B**) Fold changes in anti-insulation score by cluster for P1 to P2 (P2/P1) and P2 to P3 (P3/P2). (**C**) Odds ratio of observing P3 marker genes vs. expression-matched genes (>25th percentile of P3 marker gene expression) over matchmaker clusters (left) and grouped by matchmaker vs. not matchmaker (right). Error bars indicate the 95% confidence interval.

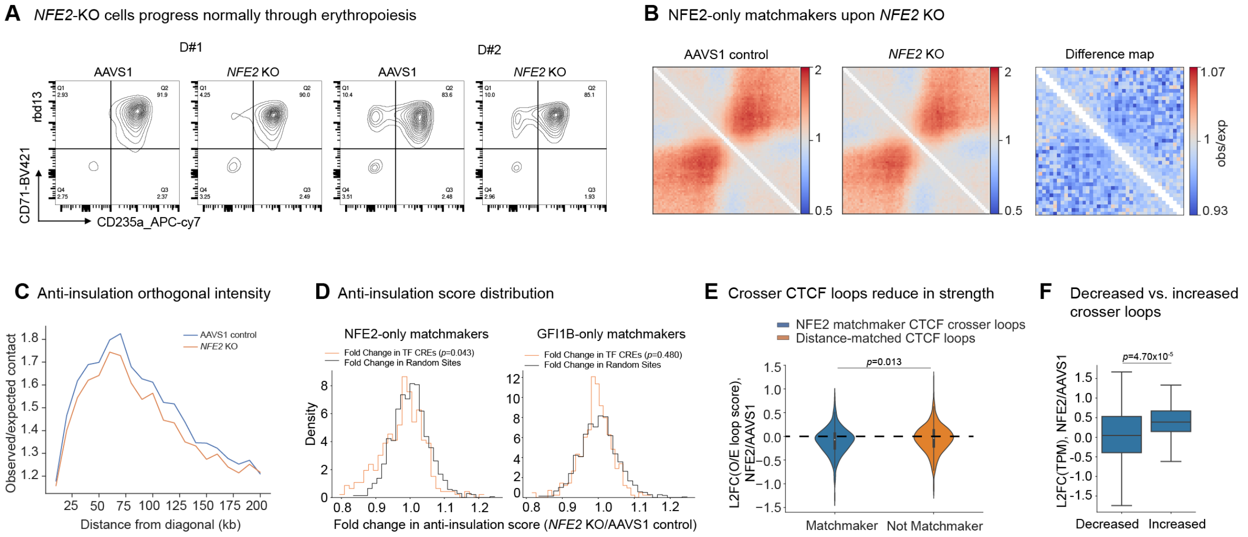

**Supplementary Figure 14: Perturbation of transcription factor NFE2.** (**A**) Flow cytometry profiles for CD235a and CD71 for the two replicates of NFE2-KO and AAVS1-targeted cell populations. Replicates represent two different biological donors. (**B**) Pileup of NFE2-only matchmakers (i.e. TF-sensitive enhancers with sensitivity only to NFE2, see Methods) in AAVS1-targeted and NFE2-KO conditions. Right panel shows a difference map (log2) at 10kb binsize. (**C**) Observed/expected contact intensity as a function of genomic distance for the 10kb bins orthogonal to the contact map diagonal, representing the contact frequency as a function of distance. (**D**) Anti-insulation score distributions at NFE2-only matchmakers (n=330), as well as at GFI1B-only matchmakers (n=337) as a control. In both, the comparison distribution is a sample of anti-insulation fold change at 1000 random sites. At NFE2-only matchmakers, p=0.043 for a difference in means between NFE2-KO and AAVS1 control anti-insulation, while p=0.480 for the same comparison in GFI1B-only matchmakers. (**E**) Observed/expected contact intensity at CTCF (E-CTCF, P-CTCF, or CTCF-CTCF) loops across NFE2-only matchmakers in the NFE2-KO Micro-C divided by the AAVS1 control Micro-C (n=253). The O/E ratio was tested (two-sided Mann-Whitney U) against the O/E ratio of distance-matched CTCF loops not near NFE2-only matchmakers (n=975; non-matchmaker loops were subsampled to 1000 and then filtered for invalid O/E scores). (**F**) Decreased and increased crosser loop scores were identified by splitting all NFE2-only matchmakers into two groups and tested with a two-sided Mann-Whitney U. Shown is the bulk transcription (TPM) at genes within 100kb of matchmakers with decreased (n=97) vs. increased (n=103) crosser loop scores.

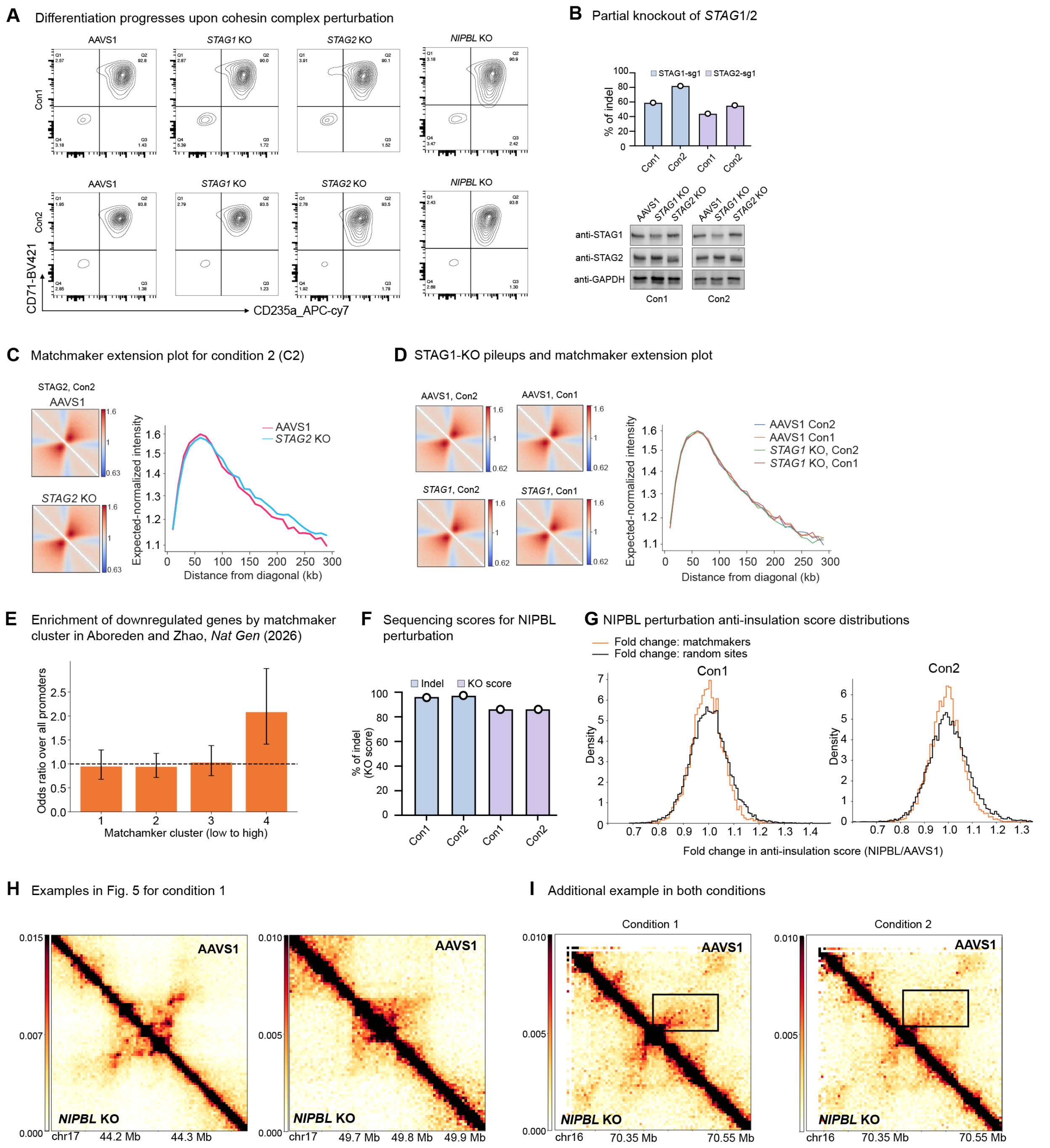

**Supplementary Figure 15: STAG1, STAG2, and NIPBL perturbations.** (**A**) Flow cytometry profiles for CD235a and CD71 for the two conditions of STAG1, STAG2, and NIPBL perturbation. (**B**) Indel rates and Western blot for STAG1 and STAG2 perturbation. Condition 1 (Con1) indicates single nucleofection at day 0, and condition 2 (Con2) indicates double nucleofection at day 0 and day 2 of erythroid differentiation. (**C**) (Left) Pileups across matchmakers in the condition not shown in **Fig. 5C**. (Right) Matchmaker length as described by the observed/expected contact frequency along the axis orthogonal to the diagonal. (**D**) STAG1 perturbation pileups and matchmaker length plots for both guide conditions. (**E**) Odds ratio enrichment near matchmakers of downregulated genes from Aboreden *et al*., 2026 (39) (Log2FC<-0.5, p-adj<0.05) upon 45 min of NIPBL depletion. Error bars indicate 95% confidence interval. (**F**) Indel rates and KO score as computed by inference of CRISPR edits (ICE) for NIPBL perturbation. Con1 indicates single nucleofection at day 0, and Con2 indicates double nucleofection at day 0 and day 2 of erythroid differentiation. (**G**) Anti-insulation score distributions for all enhancers not overlapping promoters and CTCF. (**H**) Loci shown in **Fig. 5H** for condition 1 (Con1). (**I**) Another example locus where anti-insulation is diminished in NIPBL vs. AAVS1 (chr16:70214962-70614962 at 5kb binsize).
